# Conformational state switching and pathways of chromosome dynamics in cell cycle

**DOI:** 10.1101/2019.12.20.885335

**Authors:** Xiakun Chu, Jin Wang

**Affiliations:** Department of Chemistry, State University of New York at Stony Brook, Stony Brook, New York 11794, USA; Department of Physics and Astronomy, State University of New York at Stony Brook, Stony Brook, New York 11794, USA

## Abstract

Cell cycle is a process and function of a cell with different phases essential for cell growth, proliferation, and replication. Cell cycle depends on the structure and dynamics of the underlying DNA molecule, which underpins the genome function. A microscopic structural-level understanding of how genome or its functional module chromosome performs the cell cycle in terms of large-scale conformational transformation between different phases such as the interphase and the mitotic phase is still challenging. Here, we develop a non-equilibrium excitation-relaxation energy landscape-switching model to quantify the underlying chromosome conformational transitions through (de-)condensation for a complete microscopic understanding of the cell cycle. We show that the chromosome conformational transition mechanism from the interphase to the mitotic phase follows a two-stage scenario, in good agreement with the experiments. In contrast, the mitotic exit pathways show the existence of an over-expanded chromosome that recapitulates the chromosome in the experimentally identified intermediate state at the telophase. We find the conformational pathways are heterogeneous and irreversible, as a result of the non-equilibrium dynamics of the cell cycle from both structural and kinetic perspectives. We suggest that the irreversibility is mainly due to the distinct participation of the ATP-dependent structural maintenance of chromosomal protein complexes during the cell cycle. Our findings provide crucial insights into the microscopic molecular structural and dynamical physical mechanism for the cell cycle beyond the previous more macroscopic descriptions. Our non-equilibrium landscape framework is general and applicable to study diverse non-equilibrium physical and biological processes such as active matter, differentiation/development and cancer.

## I. INTRODUCTION

The fundamental structure–function paradigm in biology is preserved for a broad range of molecules, from elementary single-molecule units such as proteins to highly complex biopolymer assemblies such as the genome.^1^ The functional 3D genome organization facilitates gene regulation by engaging two distal enhancer and promoter sequences to physical proximity.^2^ The details of the connections between the structure and the function at the genomic level are far from clear. The recently developed chromosome conformation capture techniques^3^ and the derivative Hi-C method^4^ measure the spatial proximity between genomic loci and provide static pictures of chromosomal organization at an unprecedented resolution. However, chromosome dynamics, which is closely related to the activation of transcription and gene regulation,^5,6^ remains poorly understood.

The cell cycle is a vital function of a cell, as it is crucial for cell growth, proliferation, and replication.^7,8^ The cell cycle proceeds through a few phases, notably the interphase (I phase) and the mitotic phase (M phase). The function of a cell depends on its underlying chromosomal structure and dynamics. However, gaining an understanding of the structural chromosomal transformations between the cell-cycle phases is still challenging. Hi-C data for the I phase show that a chromosome is hierarchically organized into topologically associating domains (TADs)^9–12^ and compartments.^4,13^ As the fundamental chromosomal structural units,^14,15^ TADs are characterized by having more frequent contact within these megabase-sized domains than with neighboring regions. TADs are critical for genome function at the I phase through the modulation of DNA replication^16^ and the regulation of transcription.^17^ At a higher level (>5 Mb), the compartments promote the genome function by spatially segregating the gene-rich euchromatin and gene-poor heterochromatin into a plaid Hi-C pattern at the I phase.^4,13^ Recent Hi-C experiments on the cellcycle process show that TADs and compartments vanish during mitosis.^18,19^ In addition, the experiments have indicated that there is a significant global reshaping of a chromosome from a crumpled fractal globule at the I phase^20^ to a condensed quiescent cell-type-independent cylinder at the M phase.^21^ A compressed mitotic chromosome suppresses transcription by limiting the accessibility of the promoter regions to transcription factors.^22^ On the other hand, a condensed cylinder-like chromosome favors the movement and the separation of the chromosome during cytokinesis. These features are a manifestation of the chromosomal structure–function regulation at different cell-cycle phases.

A precise description of chromosomal reorganization during the cell cycle is a prerequisite for understanding cellular function. Many studies have harnessed the power of Hi-C to measure chromosome structural at different phases during the cell cycle.^21,23–25^ However, these experiments were performed for a minimal number of cell-cycle phases, so they do not give a continuous picture of cell-cycle chromosome dynamics. Recently, time-series Hi-C experiments that measured the chromosomal reorganization at a high temporal resolution were developed and applied to the cell cycle.^19,26,27^ However, the measurements are inevitably affected by the temporal heterogeneity of different cells, so that the Hi-C data at a particular time may be from a mix of cells at different cell-cycle phases. This is a serious issue when trying to determine the contact map for a specific cell-cycle phase from time-course Hi-C data. Therefore, there is still a need for a reliable approach that can fully cover the cell-cycle spatial-temporal scale and precisely characterize chromosomal structure and dynamics during the cell cycle.

The cell cycle has high bistability in the phase transition.^28,29^ Cells at the I and M phases have distinct phenotypes and can be easily characterized in experiments. Targeted Hi-C experiments on separately arrested cells at the I and M phases can provide relatively accurate descriptions of the chromosome structures in these two phases. The chromosome dynamics at either the I or M phase is constantly influenced by the non-equilibrium effects of the surrounding cellular environment. Nevertheless, an effective equilibrium landscape can describe the non-equilibrium dynamics within each individual phase of the cell cycle in some circumstances.^30,31^ The minimally biased energy landscapes based on experimental Hi-C data using the maximum entropy principle have been successful in quantifying the spatial organization of chromosomes in the I phase^32^ and M phase.^33^ This suggests that the cell cycle can be approximately described by a combined process of intra-landscape dynamics at the individual I and M phases and inter-landscape hopping between these two phases.

However, determining the pathways and kinetics for the phase transition without sufficient time-dependent data is quite challenging. A recent computational study using a Markovian dynamics model successfully predicted the transition pathways and rates for several equilibrium and quasi-equilibrium systems. Only a few points on the transition paths were known beforehand.^34^ When only two stable states or landscapes are available, as an extreme example, an equilibrium two-basin energy landscape can be developed. The concept was initially introduced in studies of large-scale protein conformational switching.^35–39^ In practice, the landscape was constructed by inheriting properties from the individual landscapes with enforced connections at the intersections. The double-basin landscape leads to an effective intra-landscape barrier-crossing framework for the state-transition process. However, applying such methods to the cell cycle may be inaccurate. The chromosomal phase transformation during the cell cycle is driven by the extensive participation of ATP-dependent SMC complexes in chemical reactions.^40^ This indicates that the cell cycle is a far-from-equilibrium process. Thus, using an equilibrium approach may be problematic. Recently, a general approach with an optimal-transport analysis was developed for learning the continuous reprogramming trajectory from discrete time-course single-cell RNA sequencing data.^41^ The analysis focused on delineating the time evolution of out-of-equilibrium systems by capturing the time-varying probability distribution of the cells. This approach seems quite promising for investigating the chromosome dynamics of the cell cycle, since single-cell Hi-C experiments are progressing rapidly.^18,42^ However, note that the contact maps produced by single-cell Hi-C are sparse, leading to concerns about in-depth data interpretation and further structural modeling.^43^

We aim to develop a model that can simulate cell-cycle chromosome dynamics. The model should capture the non-equilibrium essence of the cell cycle while being consistent with the experimental evidence. As stated elsewhere,^44,45^ non-equilibrium dynamics can be either adiabatic or non-adiabatic, and these are associated with distinct mechanisms and kinematics. In an adiabatic regime, a non-equilibrium bistable state system has faster inter-landscape dynamics than the intra-landscape motion. The fast inter-landscape dynamics can be averaged, so the process can then be simplified so that it is effectively equivalent to a classic intra-landscape barrier-crossing model of a single effective landscape^46^ and so, analogous to equilibrium (adiabatic) dynamics. In contrast, in the non-adiabatic case, the waiting time for inter-landscape hopping is much longer than the typical timescale for intra-molecular motion. Note that the waiting time for dwelling on the intra-landscape in non-adiabatic dynamics is also much longer than the time interval of the actual inter-landscape jump. Chromosomes in the I and M phases have significantly different Hi-C patterns, so a spontaneous phase transition is highly improbable. Furthermore, the SMC complexes drive the cell-cycle chromosomal reorganization through the energy supply from ATP hydrolysis.^40^ This combination of features leads to a non-adiabatic (relatively slower inter-landscape dynamics compared to the intra-landscape dynamics) and non-equilibrium (energy supply) approximation to the cell-cycle chromosome dynamics. Notably, the process is analogous to the single-molecule motor activation associated with the large-scale conformational transition driven by the energy from ATP hydrolysis,^47^ and it is not always easy to realize adiabaticity.^48,49^

Another interesting feature of the cell cycle is the abrupt phase transformation, which occurs like a switch.^50^ A cell-cycle phase transition is governed by the underlying biochemical regulatory network, which consists of cell-cycle activators and inhibitors.^51^ Therefore, the model should include the effects on chromosome structural dynamics of the rapid change of the gene regulatory network when triggering the cell-cycle phase switch.

Here, we simulate the cell-cycle phase transition using a landscape-switching model (Fig. 1). Each transition is regarded as a combined process. There is a dwelling process in the energy landscape of one phase and then an instantaneous energy excitation that results in landscape switching followed by a relaxation process in the landscape of the new phase. The energy gained from SMC complexes by the chromosome during the cell cycle is implemented as an energy excitation and ensures the chromosome can successfully switch from one energy landscape to the other. The switching originates from the energy pump of the ATP and it breaks the detailed balance of the system, resulting in a non-adiabatic non-equilibrium process. The instantaneous energy excitation, which is regarded as a sudden stimulus that triggers the phase transition, is also consistent with the abrupt cell-cycle phase-switch concept. Therefore, the landscape-switching model captures the main characteristics of the cell cycle process and is promising, for it should result in reliable predictions. In our previous work, we demonstrated that the non-equilibrium cell-cycle dynamics is determined by both the potential landscape and the curl probability flux.^52,53^ Whereas the landscape stabilizes the states along the cell cycle, the non-zero flux, which reflects the degree of the detailed balance breaking or non-equilibrium,^54^ drives the stable cell-cycle oscillations. The non-equilibrium effects in the landscape-switching model are the driving forces for the cell-cycle progression and are reminiscent of the curl flux in our previous theoretical studies.

**FIG. 1.**
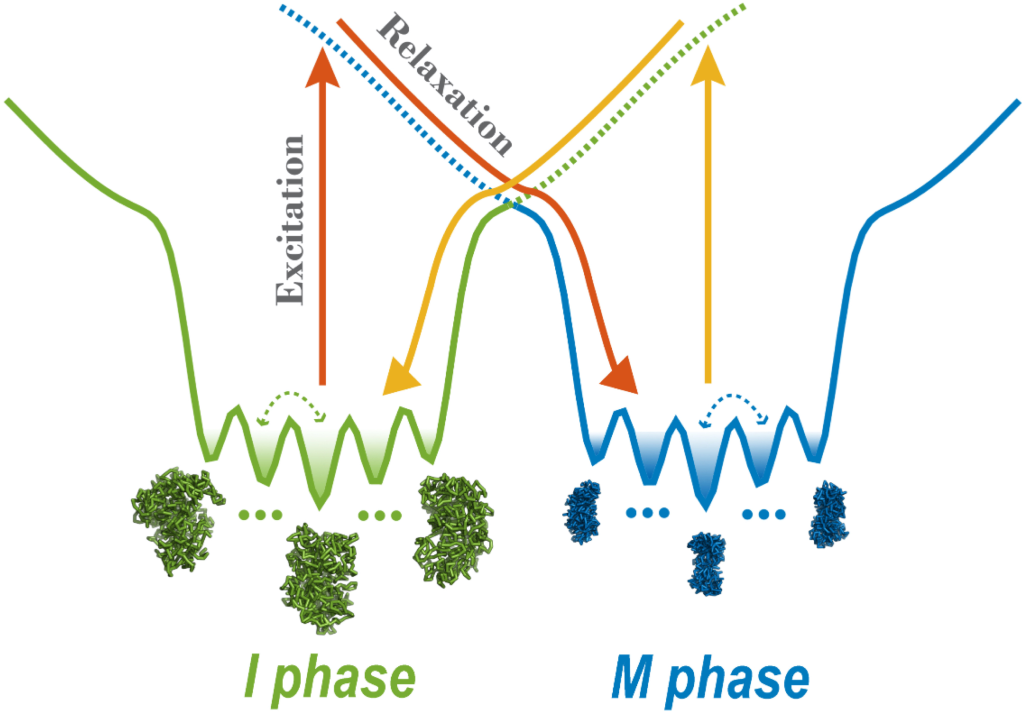
Energy landscape-switching model for simulating chromosome conformational transition dynamics during the cell cycle. There are two energy landscapes, each with a prominent basin, representing a chromosome in the I and M phase, respectively. Within each basin, there are many potential-energy minima, corresponding to different metastable chromosome states. The simulation starts with the chromosome structure in the I or M phase and runs for a period of time. Then, a sudden excitation due to switching the energy landscape to the M or I phase is implemented. Finally, the system relaxes to the new basin in the post-switching energy landscape.

Using this landscape-switching model, we identified a statistically significant number of pathways for the chromosome that are dependent on cell-cycle dynamics. These are irreversible and highly heterogeneous, though not random. Our results are in good agreement with the contemporary experimental characterization of the mechanisms for mitosis and the mitotic exit process at the chromosomal level.^18,19,21,26^ We found that the formation of SMC complex-mediated loops is irreversible during the cell cycle, giving rise to a biological explanation for the irreversibility from a structural perspective. From a physical perspective, the irreversibility uncovered by our model is controlled by the broken detailed balance of the underlying non-equilibrium process, often in the form of extended ATP hydrolysis for energy pumping. The physical mechanism we obtained provides a quantitative basis for deciphering the dynamic regulation of the chromosome structure–function relation during the cell cycle. The model has potential applications for a wide range of biological processes, which are mostly non-equilibrium.

## II. METHODS

### A. Hi-C data processing

The Hi-C data for the I and M phases for HeLa cells were downloaded from the ArrayExpress database with the accession number E-MTAB-1948.^21^ The Hi-C data of different sub-stages within the I phase display quite similar patterns at both short- and long-range pairwise contact regions and are highly correlated.^21^ Therefore, we used mid-G_1_ Hi-C data to represent approximately the entire I phase. On the other hand, we used metaphase Hi-C data to represent the M phase. Two replicas of the Hi-C data from each phase were combined together, and the analysis proceeded routinely using the HiC-Pro software.^55^ We normalized the pairwise contact frequencies into contact probabilities, which are easier to implement in the following polymer simulations, based on the reasonable assumption that neighboring beads are always in contact with a probability *P*_*i,i* ±1_ ≡ 1.0.^32^ All the Hi-C matrices had 100-kb resolution, which is high enough to monitor the chromosome dynamics. We focused on a long segment on chromosome 5 from 80 to 161.1 Mb, which eventually gave 812 beads in the system in the following polymer simulations.

### B. Coarse-grained chromosome model

A generic bead–spring polymer model was used as the basis and background to simulate the chromosome dynamics.^56^ The potential of the generic polymer model *U*_Polymer_ was constituted from the traditional bond, angle, and non-bonded interaction potentials, as explained in the following. Neighboring beads (*i, i* + 1) were connected by pseudo-bonds through the finitely extensible nonlinear elastic (FENE) potential.^57^ To avoid any numerical instability due to divergence of the FENE potential at ∼*R*_0_, we use a method proposed by Tokuda *et al*.,^58^ in which the bond potential *V*_Bonds_ switches from the FENE potential to a power-law function near *R*_0_:

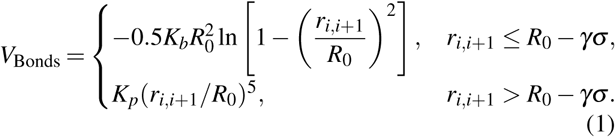

To avoid an unrealistic overlap between neighboring bonded beads, a hard-core repulsive potential, derived from the Lennard-Jones potential, was added:

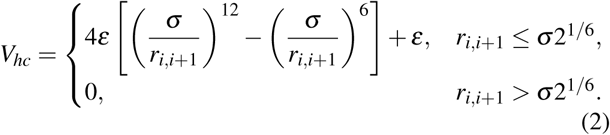

The angle potential was added for every three adjacent beads (*i* − 1, *i, i* + 1)^59^:

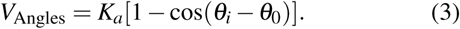

The effects of topoisomerases, which help to relieve the formation of knots, were mimicked by allowing chain-crossing in the model.^21^ This was done by implementing a soft-core repulsive potential between all the non-bonded pairs (*i, j*)^32^:

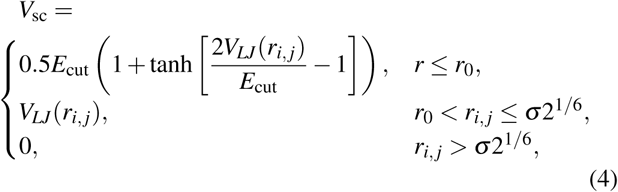

where

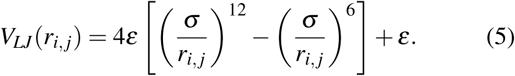

A spherical confinement with the potential *V*_*C*_, which has a semi-harmonic potential, was used to mimic the volume fraction of the chromosome:

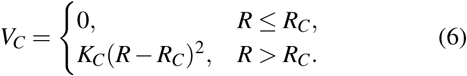

Any of the beads in the polymer are pulled back if their distance to the center of the simulation box is more than *R*_*C*_.

The simulations were performed by Gromacs (version 4.5.7)^60^ with the PLUMED plugin (version 2.5.0).^61^ Reduced units were used. The energy unit was *ε* = 1.0. The bond length σ was set to be the length unit. The parameters used for the FENE potential in the simulations have been widely used elsewhere.^58,59^ *R*_0_ = 1.5σ allows the bonds to stretch flexibly and *K*_*b*_ = 30.0*/*σ ^2^. The power-law potential in *V*_Bonds_ has *K*_*p*_ = 100.0 and *γ* = 0.125 to ensure there is a smooth connection near *R*_0_ (for both the potential and the force). The angle potential has a strength of *K*_*a*_ = 2.0 and angle of *θ*_0_ = *π*.^59^ In the soft-core potential, *E*_cut_ = 4.0, we set

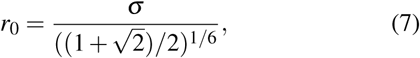

so *U*_*LJ*_(*r*_0_) = 0.5*E*_cut_ = 2.0.^32^ To ensure the volume fraction of the chromosome at 10% was the same as used previously,^32,59^ we set *K*_*C*_ = 100.0*/*σ ^2^ and *R*_*C*_ = 9.0σ in the confinement potential. The temperature in all simulations was set to 1.0 in energy units by multiplying by the Boltzmann constant. Langevin stochastic dynamics was applied with a time step of 0.001*τ* and a friction coefficient of 1.0*τ*^−1^, where *τ* is the reduced time unit.

### C. Chromosome simulations guided by maximum entropy principle

The chromosome ensembles in the I and M phases were produced using the maximum entropy principle approach by practically incorporating the experimental Hi-C data into the generic polymer simulations. Based on the experimental data (i.e., Hi-C data), a new energy function can be written:

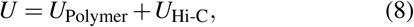

where *U*_Polymer_ is the generic polymer potential described above and has the following expression:

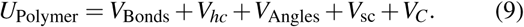

Here, *U*_Hi-C_ is the experimental Hi-C restraint potential and should have a specific functional form that is linear for the pairwise contact probabilities, following the maximum entropy principle.^62^ Also, *U*_Hi-C_ is the sum of the pairwise non-bonded contact probabilities, which corresponds directly to the Hi-C data map:

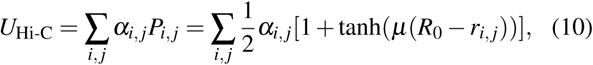

where the step function *P*_*i, j*_ represents the contact probability between the genomic locus *i* and *j. α*_*i, j*_ is iteratively adjusted to match *P*_*i, j*_ to the experimental Hi-C data *f*_*i, j*_ via the maximum entropy principle approach. We set *µ* = 3.0 in the expression for *P*_*i, j*_. The details of the iteration procedure can be found in Refs. 32 and 33.

The main outcome of the maximum entropy approach is that it generates an ensemble of chromosome structures that collectively reconstruct the experimental Hi-C data while remaining as close as possible to the prior distribution from the generic polymer simulations through maximizing the relative entropy.^62^ More importantly, the maximum entropy principle can give rise to an interaction potential energy function (i.e., force field) for simulations that are rationally restrained by the experimental ensemble-averaged Hi-C data, and this eventually leads to a minimally biased energy landscape.^32^ Such an effective energy landscape not only can give an accurate description of chromosome thermodynamic distributions inferred from the Hi-C data-based contact probabilities, but it can also provide the details of the kinetics within one phase during the simulations (Fig. S6). Therefore, simulations based on the maximum entropy principle eventually lead to two potentials, *U* (***r***|*I*) and *U* (***r***|*M*), which, respectively, describe the chromosome conformational dynamics for the I and M phases. Here ***r*** is the coordinate of the system. Note that the only difference between *U* (***r***|*I*) and *U* (***r***|*M*) is the non-bonded experimental Hi-C restraint term *U*_Hi-C_, in particular at *α*_*i, j*_.

### D. Cell-cycle dynamics revealed by energy landscape-switching model

We developed an energy landscape-switching model to simulate the chromosome transformation during the cell cycle. The model goes beyond the previous studies based on allosteric protein dynamics.^63,64^ The cell-cycle process has been simplified to two transitions: I → M and M → I. The simulation algorithm had the following three steps. First, the chromosome in the I or M phase was simulated by the corresponding potential energy landscape *U* (***r***|*I*) or *U* (***r***|*M*), obtained from the maximum entropy simulation for a duration of 5 *×* 10^3^*τ*. Then, the potential landscape was switched from *U* (***r***|*I*) to *U* (***r***|*M*) for the I → M transition or from *U* (***r***|*M*) to *U* (***r***|*I*) for the M → I transition. This is reminiscent of the energy excitation process due to the chemical reactions of the SMC complexes pumped by the underlying extended forms of ATP hydrolysis in real cells.^40^ This implementation represents the energy pump through ATP hydrolysis that breaks the detailed balance of the system, resulting in a non-equilibrium process. Finally, the simulations under the new energy landscape of the M phase with *U* (***r***|*M*) or the I phase with *U* (***r***|*I*) were run for 1 *×* 10^4^*τ*. For each trajectory, the waiting time for dwelling at one phase was 5 *×* 10^3^*τ*, which corresponds to the time of an inter-landscape hopping event. We calculated the decay of the autocorrelation function of the chromosome order parameter^65^ and estimated the timescale for intra-landscape dynamics at one phase (Fig. S1). As illustrated in Fig. S1, although the actual inter-landscape excitation is instantaneous, the inter-landscape hopping event occurs over a much longer timescale than that for the intra-landscape dynamics. These features lead to the non-adiabatic non-equilibrium dynamics, which we simulated.

In practice, the simulations were initialized from the structures obtained after clustering the trajectories of the maximum entropy principle for the I or M phase, and only the clusters with populations higher than 1% were selected. There were 52 and 56 clusters for the I and M phase chromosome ensembles, respectively. For each cluster, the five structures closest to the cluster center were chosen. Therefore, we accumulated 260 transitions from the I to the M phase, and 280 transitions from the M to the I phase.

## III. RESULTS

### A. Cell-cycle chromosome conformational dynamics triggered by energy landscape switching

We use a generic polymer model restrained by the Hi-C data through the maximum entropy principle to simulate the chromosome conformational dynamics in the I and M phases individually (Section II). The models successfully reproduced the experimental Hi-C data for the I and M phases (Figs. S2 and S3). Besides, we observed formations of crumpled fractal globular chromosomes in the I phase and condensed cylindrical chromosomes in the M phase (Figs. S4 and S5). These results confirm the validity of our approach to modeling the chromosome structural ensemble using Hi-C data. Furthermore, we investigated the spatiotemporal chromosome dynamics in the I phase by calculating the mean squared displacement of the chromosomal loci (Fig. S6). We observed that there was sub-diffusion with scaling factor *β* = 0.45 in mean squared displacement ∼*t*^*β*^ in the I phase, like that found in the experiments (*β* ≈ 0.39–0.44).^66^ Therefore, the potentials that represent effective energy landscapes can successfully capture the correct thermodynamics and kinetics of the chromosome dynamics in the individual phases. These results provided the basis for the following landscape-switching simulations.

The cell-cycle process in the simulations is divided into the I → M and M → I phase-to-phase transitions. We simulated each phase transition by a combined process based on the initial dwelling dynamics at the I (M) phase, then instantaneous landscape switching, and finally the relaxation dynamics at the M (I) phase (Fig. 1; Section II). We observed significant chromosome conformational changes during the cell-cycle simulations. The circle at the center of Fig. 2 illustrates the apparent cyclic change of the contact probability *P*_*s*_ for the genomic distance *s* during the cell cycle.^20^ *P*_*s*_ characterizes the chromosome conformational transition between the fast decrease *s*^−1^ in the I phase and the slow decrease *s*^−0.5^ in the M phase (Fig. S10). As the chromosomes may be trapped in multiple disconnected metastable states for a remarkably long time in either the I or M phase (Figs. S6 and S7),^67^ the energy landscape-switching simulation has efficiently activated the cell-cycle dynamics. Therefore, the simulations require only reasonable computational costs.

**FIG. 2.**
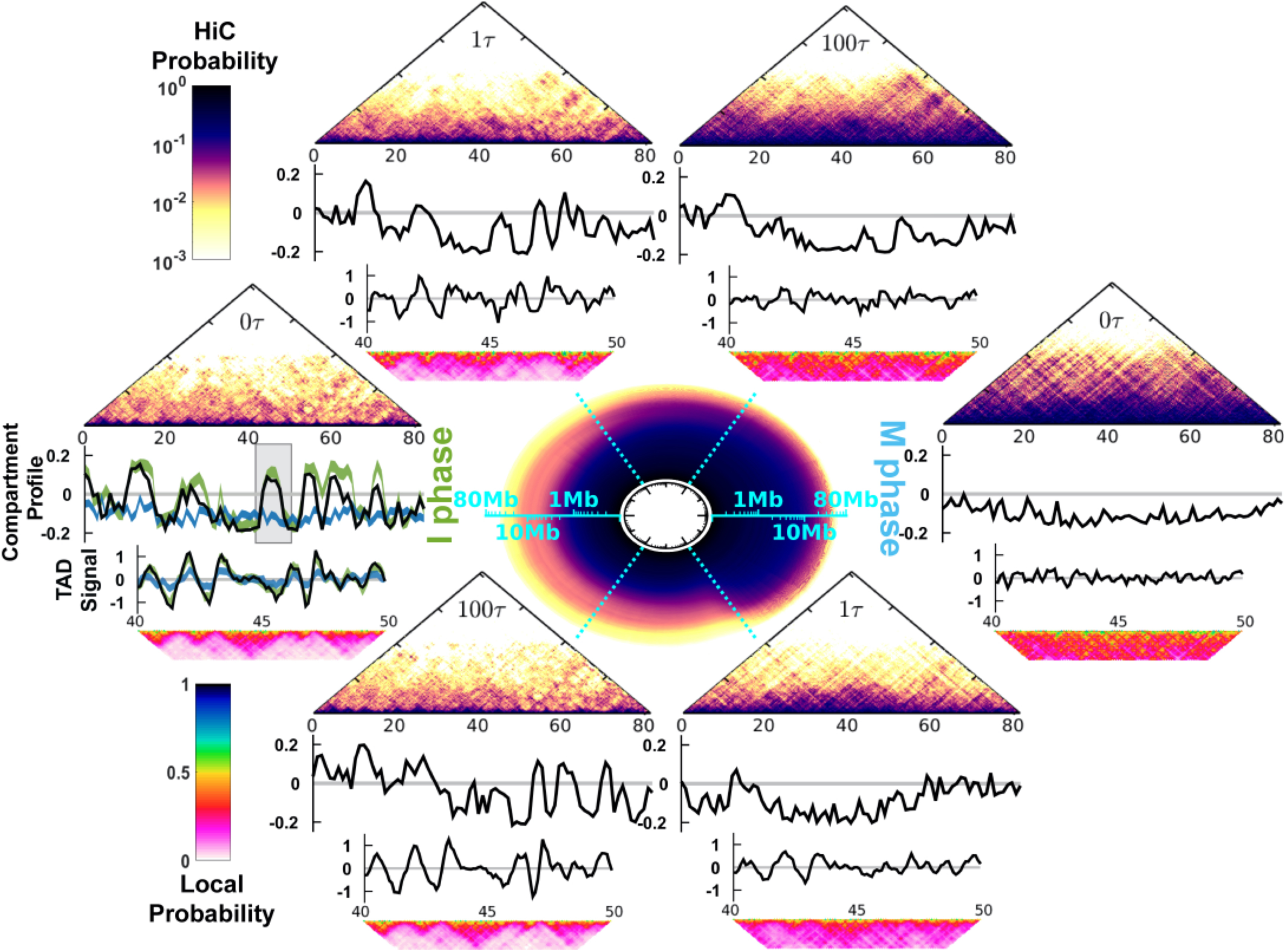
Chromosome conformational transitions during the cell cycle. The circle at the center represents the contact probability *P*_*s*_ for the genomic distance *s*, which evolves during the clockwise cell cycle. The radial and angular coordinates of the circle are, respectively, the genomic distance *s* and cell-cycle processing time *t*, both shown in logarithmic scale. The upper and lower semicircles describe the transition from the I to the M phase and from the M to the I phase, respectively. At particular time points *t* = 0*τ*, 1*τ*, and 100*τ*, the corresponding Hi-C contact map, compartment profile, TAD signal, and short-range contact map for the 40–50 Mb region are shown. At the far left, the green and blue lines in the compartment profile and TAD signal plots are the values calculated from the Hi-C data for the I and M phases, respectively. The shaded region in the compartment profile indicates the chromosome segment (40–50 Mb) for the TAD signal and short-range contact map plots.

The evolution of the contact probability map shows that the chromosome conformational transition for the I → M transition (M → I transition) proceeds with condensation (de-condensation) followed by the loss (formation) of the TADs and the compartments (Fig. 2; see the Supplementary Information for the details of the calculation of the TAD signal and the compartment profile). During the transition from the I to M phase, the chromosomes initially at *t* = 0*τ* exhibit the characteristics of the I phase. After a very short time lag to *t* = 1*τ*, there are notable widespread changes in the contacts (Fig. S15), leading to deviations of the TAD and compartment profiles from those in the I phase. Further progression of the cell cycle continuously increases the extent of condensation of the chromosome along with the gradual loss of the TADs and the compartments. The TADs and the compartments have almost disappeared at *t* = 100*τ*. Further analysis of the kinetics of the TAD signals showed that there was a very fast deformation of the TADs (Fig. S11). Such a rapid and acute reorganization of the chromosome structure during phase switching was also found experimentally when G_2_ cells enter the prophase.^19^

In the M → I transition, we observed that the TADs rapidly form very early at *t* = 2*τ* (Fig. S11). On the other hand, the compartment profile at *t* = 10*τ* is still very different from that in the I phase (Fig. S13). Therefore, the compartments form more slowly than the TADs. The asynchronous establishment of the TADs and the compartments was observed experimentally in the time series Hi-C data measured for the mitotic exit process.^26^ The experimental analysis concluded that the TADs rapidly form before the entry into the G_1_ phase, while compartmentalization is slow and still proceeds after the G_1_ phase.^26^ Note that more contacts can be lost at *t* = 1*τ* than at the final destined I phase (Fig. S14). This observation implies that a chromosome may have an over-expanded conformation around *t* = 1*τ* during the M → I transition.

Overall, we showed that a chromosome undergoes condensation and de-condensation associated with deforming and forming of the TADs and compartments during the cell cycle. The chromosome conformational transitions in both the I → M and M → I processes occur very fast. This may be due to the instantaneous switching model, which produced strong driving forces to guide the motion of the system in the post-switching landscape. There is recent and increasing experimental evidence to show that the chromosome undergoes rapid structural reorganization during the cell-cycle phase transition,^19,26,27,68^ which occurs abruptly in cells. A sudden change of the external environment during phase switching can dramatically induce strong conformational strains on the chromosome, so this requires a fast structural reorganization.

### B. Chromosome conformational evolution during the cell cycle

To quantify the chromosome conformational transition pathways, we collected all the trajectories and projected them onto several order parameters (Figs. 3 and 4). The phase-transition trajectories readily reached the destined phase during the simulations, confirming that our simulations had converged (Figs. S16 and S17). First, we focused on the global shape change of the chromosome conformation by examining the lengths along the principal axes (PAs),^33^ aspherical shape parameter (Δ),^69^ and radius of gyration (*R*_*g*_) (see the Supplementary Material for definitions).

**FIG. 3.**
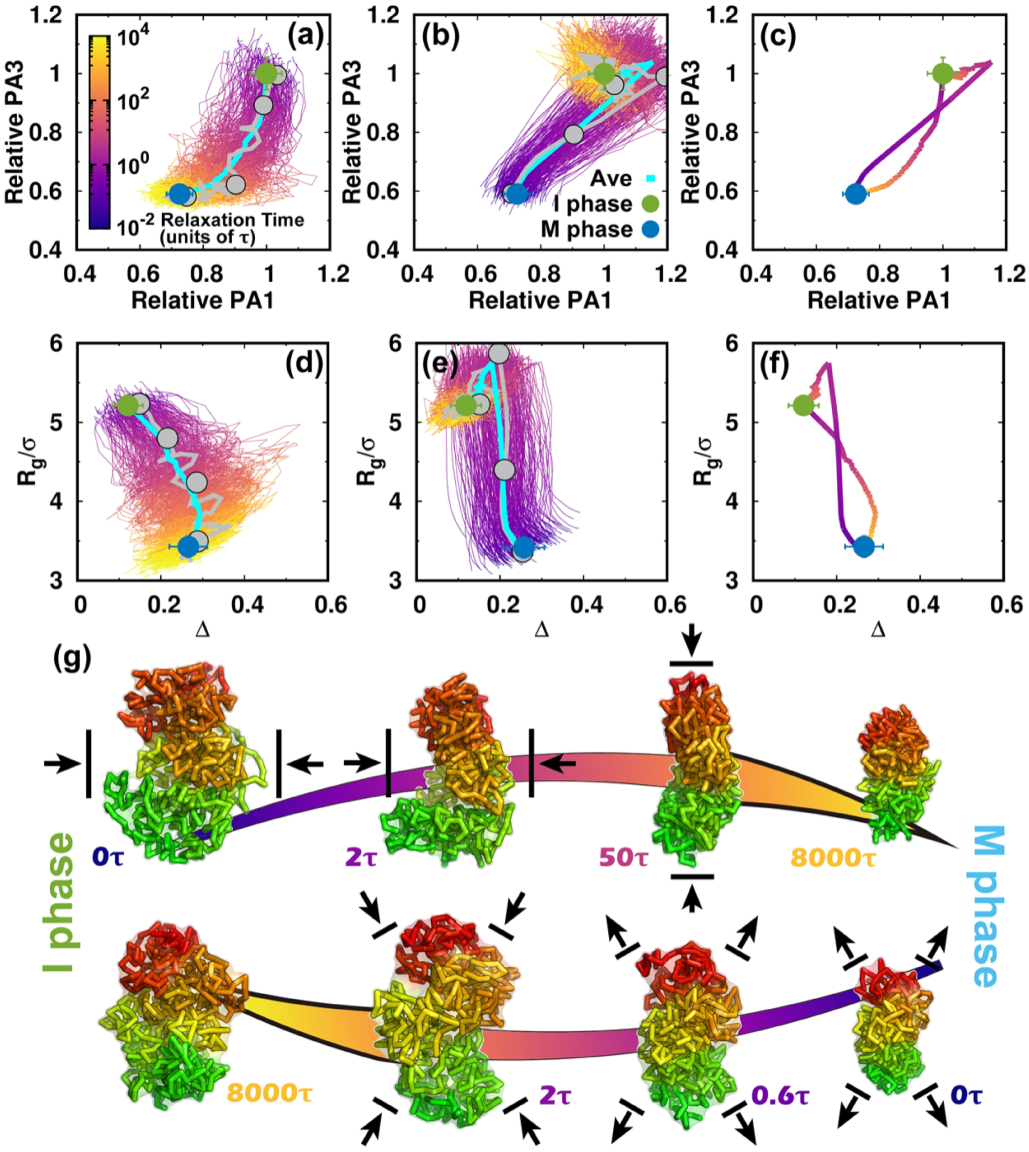
Shape changes of the chromosome during the cell cycle. (a) The pathways projected onto the first and third principal axes (PAs) during the transition from the I to the M phase. The first and third PAs are defined to be the longest and shortest in the chromosome conformation, respectively. The PA lengths are relative to the mean values of those at the I phase. Thus, the transition approximately starts from (1, 1) and decreases during the condensation process to the M phase. All the transition trajectories are shown and colored by time in a logarithmic scale. The green and blue dots represent the mean values for the I and M phases. The error bars correspond to the standard deviations. The pathway averaged by all trajectories is shown with the cyan line. (b) Like (a) but for the transition from the M to the I phase. (c) The average pathways colored using a logarithmic timescale for the two directional transitions. (d) The pathways projected onto the aspherical parameter Δ and radius of gyration *R*_*g*_ during the transition from the I to the M phase. Δ is in the range 0–1. For perfect spheres, Δ = 0. (e) Like (d) but for the transition from the M to the I phase. (f) The average pathways for the two directional transitions. (g) Typical chromosome structures extracted from the transition trajectories. Upper: I → M transition. Lower: M → I transition. The pathways for the selected structures are shown in (a), (b), (d), (e) by gray lines with dots corresponding to the positions of the structures. The chromosome structures are colored from red to green based on the genomic distance. The black arrows for the chromosome structures illustrate the major shape changes expected in the next step.

**FIG. 4.**
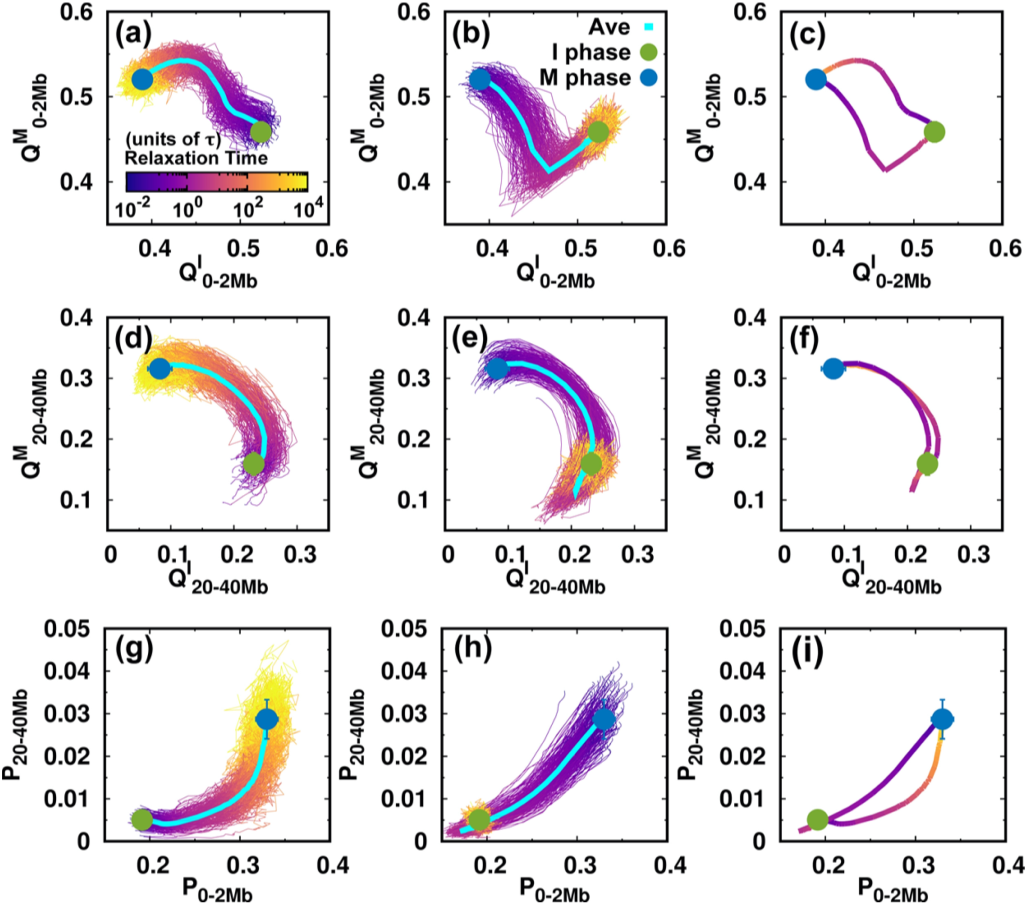
Contact changes of the chromosome during the cell cycle. (a–c) The pathways projected onto the local conformational order parameters. 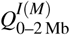 is the fraction of contacts formed by the pairwise loci that are in the genomic distance from 0 to 2 Mb, with superscript I and M, respectively, corresponding to the references chosen from the I and M phases. The references are the average pairwise distances of the I or M phase ensembles (Figs. S4 and S5). Pathways for (a) the I → M transition, (b) the M → I transition, as well as (c) the average for the two directional transitions. (d–f), (g–i) As (a–c) but projected onto the non-local conformational order parameter 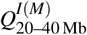 and contact probability *P* at the short (0–2 Mb) and long-range genomic distances (20–40 Mb), respectively.

Once energy landscape-switching has been triggered for the I → M phase transition, the chromosome starts to become compacted. The longest principal axis (PA1) and shortest principal axis (PA3) then decrease (Fig. 3(a)). The individual pathways are highly heterogeneous, and no dominant path can be defined. Nevertheless, all the stochastic pathways follow particular routes for condensation. We can readily see that most of the chromosomes are more compact in terms of PA3 than PA1 in the early stage. Shortening of PA1 becomes more significant for most trajectories in the late stage. Further projecting the pathways onto *R*_*g*_ and Δ gives a consistent and complementary explanation for the transition (Fig. 3(d)). In the beginning, *R*_*g*_ and Δ decrease and increase, respectively, and nearly synchronously, resulting in an anisotropic compaction of the chromosome conformation. A decrease of *R*_*g*_ along with no change or even a slight decrease of Δ occurs in the late stage, which is consistent with the idea that the longest PA1 shortens to form the mitotic conformation at the end of the transition. The typical structures extracted from one particular trajectory, which was close to the average pathway, clearly show how the chromosome evolved into a condensed cylinder (Fig. 3(g)). This is in good agreement with the two-stage mitotic chromosome condensation hypothesis based on experimental findings, in which the interphase chromosome linearly compacts into consecutive loops followed by axial compression.^21^

Interestingly, the M → I transition uses a different mechanism for chromosome de-condensation, as it is not a simple reversal of the chromosome condensation in the I → M transition. As shown in the trajectories projected onto the PAs (Fig. 3(b)), the chromosomes expand rapidly and isotropically in the early stage. Then the chromosomes preferentially reach a distinct set of conformations that have an even longer PA1 than in the I phase. This can also be seen by looking at the pathways projected onto *R*_*g*_ and Δ (Fig. 3(e)). *R*_*g*_ continues to increase in the early stage until it is bigger than in the I phase (Fig. 3(e)). This indicates that the conformation over-expands, primarily along its longest axis. The most significant chromosome conformational expansion occurred at ∼2*τ*, when *P*_*s*_ was decreasing rapidly as *s*^−1^ (Fig. S10(c)), as in the I phase. The quenching from the over-expanded chromosome conformation to that at the I phase requires compaction of the PA1 dimension, resulting in condensation to the globule-like shape. This is seen as a simultaneous decrease of both *R*_*g*_ and Δ. Finally, the average pathways for the two directional transitions connecting the I and M phases are completely different (Figs. 3(c) and 3(f)). These observations indicate that the cell-cycle dynamics of chromosome conformational switching is stochastic and irreversible.

We then used the parameters related to contact formation to project the pathways (Fig. 4). In the I → M transition (Figs. 4(a) and 4(d)), we see that local and non-local chromosome conformations have an asynchronous step in conformational switching. This can be observed by retrieving the pathways projected onto the local and non-local contact probabilities (Fig. 4(g)). The local contacts tend to form before the non-local ones in the transition from the I to M phase. On the other hand, formation of a local contact proceeds along completely different pathways during the M → I transition. We observe that 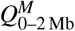 decreases considerably until it is even lower than in the final I phase. Such over-expansion can also be observed for the non-local contacts (Fig. 4(e)) and by the pathways projected onto the contact probabilities (Fig. 4(h)). Therefore, we have shown that the over-expansion of the chromosome conformation in the M → I transition occurs universally throughout the genomic sequence.

The irreversibility of the cell-cycle dynamics of the chromosome conformational transition is clearly shown by the average pathways for the two directional transitions (Figs. 4(c), 4(f), and 4(i)). The formation of local contacts is along two completely separate non-overlapping pathways for the two transitions that constitute the cell cycle. When the range of the genomic distance of the pairwise contact increases until it becomes non-local, the pathways tend to coin-cide, in particular in the stage close to the M phase (Figs. 4(f) and S18). Note that the TADs form at the local range (∼2 Mb) (Fig. S12), so that the formation and deformation of the TADs during the cell cycle follow totally irreversible pathways.

### C. Heterogeneity of TAD conformational transition during the cell cycle

To see how TADs evolve during the cell cycle, we first detected 40 TADs on the chromosome that we were investigating using the software HiCSeg.^70^ Then we calculated *R*_*g*_ for each TAD for all the trajectories and we monitored the average pathways during the transitions. In the I → M transition, most of the TADs condensed, though already abundant contacts had formed within the TADs.^10^ There were roughly two classes of *R*_*g*_ evolution: one in which *R*_*g*_ decreases monotonically to that at the M phase and the other in which *R*_*g*_ increases at first and then decreases (Fig. 5(a)). The latter resembles the results from the recent single-cell Hi-C experiments, in which some TADs underwent de-condensation from the early G_1_ to the S phase before the completion of the I phase.^18^ For each class, two typical TADs were chosen. For both TADs, the variances of the pathways were large, indicating that even one TAD can follow highly heterogeneous pathways to realize the transition.

**FIG. 5.**
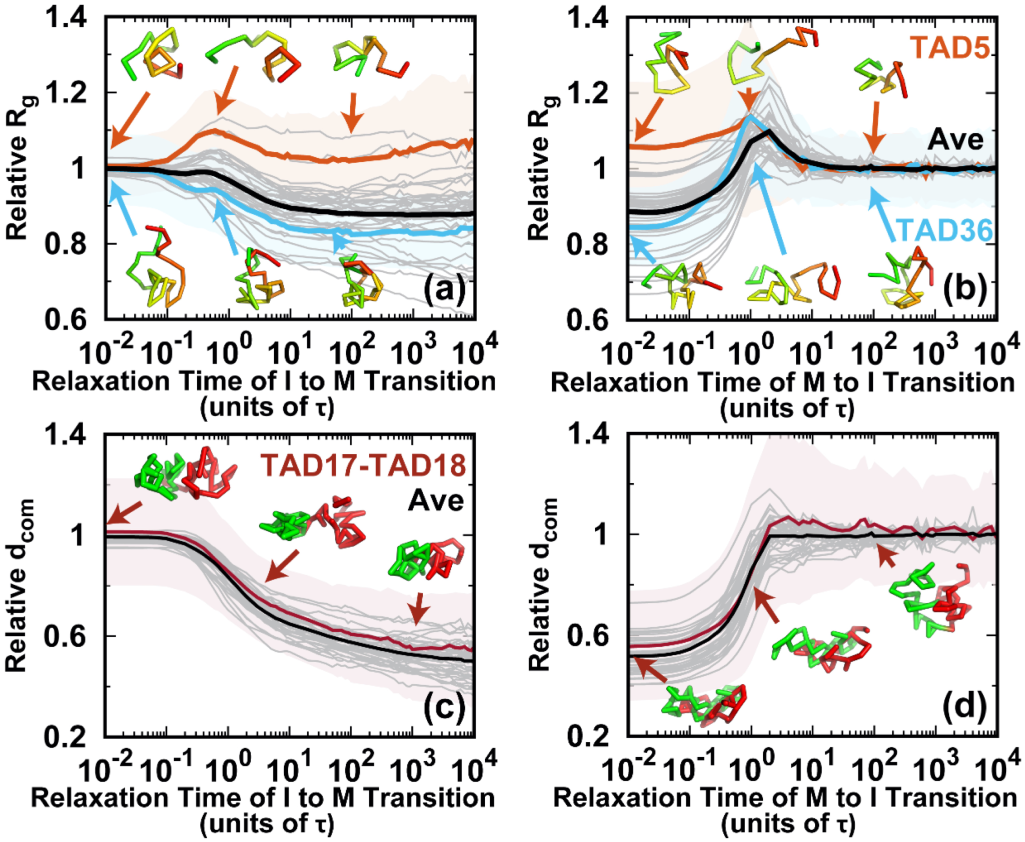
Conformational dynamics of the TADs during the cell cycle. The pathways for the radius of gyration *R*_*g*_ of the TADs for the (a) I → M and (b) M → I transitions. *R*_*g*_ relative to the mean value in the I phase for all 40 TADs is shown as gray lines. TAD5 (fifth TAD, orange, the same notation used thereafter) and TAD36 (blue) are selected for illustration. The shaded regions are the corresponding variances. Representative structures of the selected TADs are shown and colored from red to green based on genomic distance. The pathways for the distance of the center of mass (*d*_com_) between neighboring TADs for the (c) I → M and (d) M → I transitions. *d*_com_ relative to the mean value in the I phase for all 40 TADs is shown as gray lines. The distance between TAD17 and TAD18 (TAD17-TAD18) is selected for illustration. The shaded regions are the variances. The representative structures TAD17 and TAD18 are colored red and green, respectively.

On the other hand, in the M → I transition, almost all the TADs undergo de-condensation before the final condensation. However, the magnitude of the increase of *R*_*g*_ was different. The initial de-condensation may be attributed to the over-expansion, as can be observed in Figs. 3 and 4. The decondensation for all TADs reached a peak at around 2*τ*, when the chromosomes formed the over-expanded conformations. These results imply that the conformational dynamics of the TADs are cooperative processes in the M → I transition.

One of the prominent characteristics of the M phase is the loss of the TAD signals, as there is no TAD boundary due to the high condensation.^21^ The strength of the TAD signal can be measured by the amount of insulation at the TAD boundary,^71^ which is defined through the degree of the contact depletion crossing the TAD borders.^18^ The structural evolution of the TAD boundary was monitored by calculating the spatial distance *d*_com_ between two neighboring TADs during the transitions (Figs. 5(c) and 5(d)). A short (long) *d*_com_ corresponds to weak (strong) TAD insulation and weak (strong) formation of the TAD boundary. Although there were still highly heterogeneous pathways for one single inter-TAD distance evolution, the spatial distances of the average pathways all monotonically decrease to those at the M phase in the I → M transition. On the other hand, the over-expansion in the transition from the M to the I phase appears to be minor, and most of the spatial distances of the pathways increase monotonically until they reach a plateau. Therefore, the inter-TAD distance evolves more cooperatively than the intra-TAD does and it may share the same routes in the two directional transitions.

### D. Chromosome conformational kinetics during the cell cycle

We monitored the evolution of the contact probability at each locus as it interacted with others at different genomic distances and calculated the half-lives during the two directional transitions of the cell cycle (Fig. 6). In the I → M transition, the highly heterogeneous evolution of the short-range contact is shown as the extent of pathway heterogeneity weakening as the genomic distance increases (Figs. 6(a), 6(b), and S21(a)). There is a positive monotonic relation between the mean *τ*_1*/*2_ and the genomic distance (Fig. 6(c)). In particular, the local (0–2 Mb) conformations form much faster than the other non-local ones do. However, by carefully examining the second-order movements of *τ*_1*/*2_, we observe a significant fluctuation of *τ*_1*/*2_ at the local genomic distance of 0–2 Mb. This strongly suggests that the conformational kinetics for each locus is a non-Poisson process. The quantity *τ*_1*/*2_ is not self-averaging,^72^ as the distribution has a long tail far from the mean value (Fig. S21(b)). Since the formation of the TADs occurs for a genomic distance of ∼2 Mb (Fig. S12), we suggest that the TAD conformational dynamics in the I → M transition is highly heterogeneous for an individual locus. Thus, the TADs may form quite late, though the average conformational transition pathways appear to be efficient. When the genomic distance increases from 0–2 Mb to 2–5 Mb, the second-order movement of *τ*_1*/*2_ decreases sharply to near 2 and keeps decreasing smoothly for longer genomic distances. This indicates that at genomic distances longer than the TAD range, chromosome conformational switching is highly cooperative among each locus and proceeds more slowly than at the TADs.

**FIG. 6.**
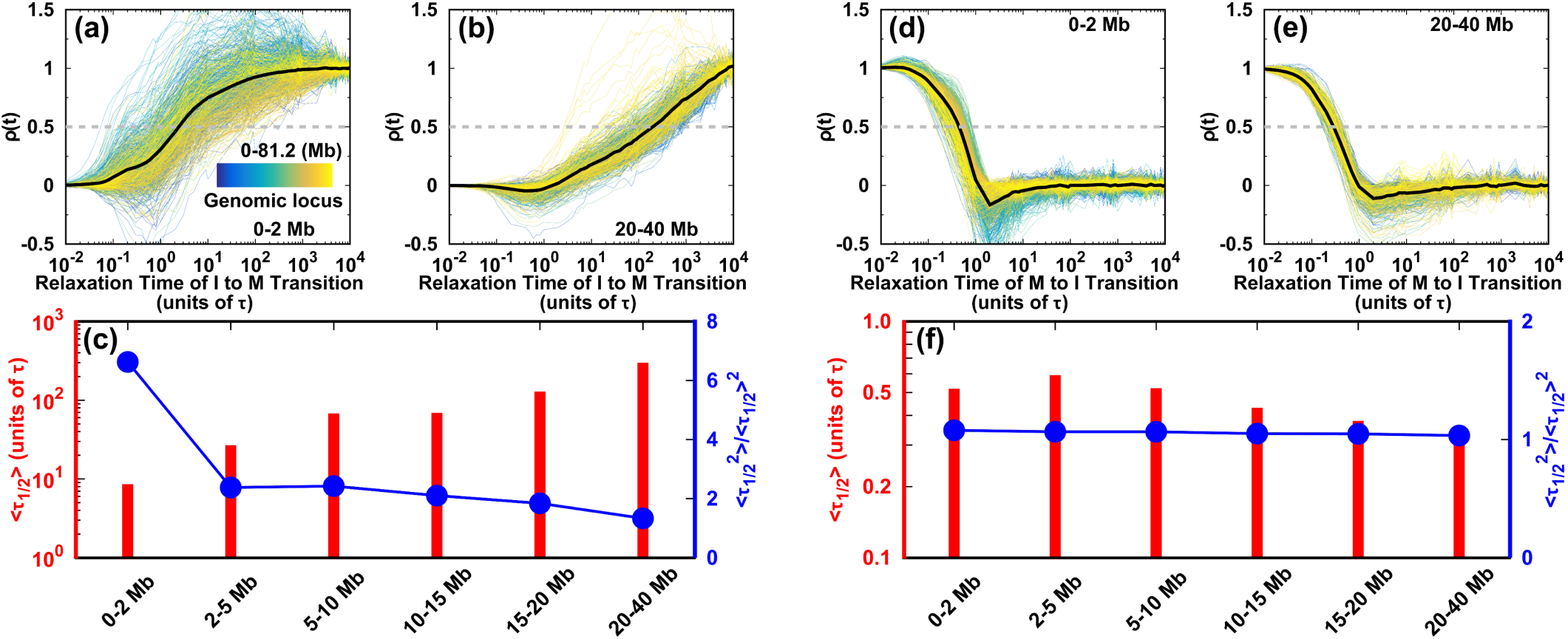
Kinetics of the chromosome conformational transition during the cell cycle. The degree of contact formation is calculated as *ρ*(*t*) = (*P*(*t*) − *P*^*I*^)*/*(*P*^*M*^ − *P*^*I*^). The transition pathways for all the individual loci are shown. (a), (b) *ρ*(*t*) for short- and long-range contacts in the I → M transition, respectively. (d), (e) As (a) and (b), but for the M → I transition. The gray dashed lines in (a), (b), (d), and (e) show where *ρ*(*t*) is half its maximum value. The associated times are the half-lives *t*1*/*2. (c) *t*1*/*2 and its second-order moment for contacts forming at different ranges during the transition from the I to M phase. (f) As (c) but for the transition from the M to I phase.

On the other hand, there appears to be little difference in the conformational evolution of an individual locus at various genomic distances during the transition from the M to the I phase (Figs. 6(d)–6(f) and S21(c)). For different contacting ranges, the mean *τ*_1*/*2_ has very similar small values, and the second-order movements of *τ*_1*/*2_ are relatively low close to 1, resulting in similar *τ*_1*/*2_ distributions (Fig. S21(d)). These observations suggest that each individual chromosomal locus acts cooperatively in the conformational switching from the M to the I phase. The kinetic rates in the M → I transition are much faster than those in the I → M transition, underlining the kinetic irreversibility of the cell-cycle dynamics.

### E. Identification of bimodal structural transition state ensemble

Precisely characterizing the transition state (TS) ensemble has always been challenging in molecular dynamics simulations. It is expected to be even more difficult in a landscape-switching simulation, as the dynamics is non-adiabatic. Nevertheless, a simulation can approximately be performed using the structural reaction coordinates and assuming the TS structurally forms halfway during the transition.^64^ This assumption resembles the widely used kinetic TS definition in conventional molecular dynamics, in which the TS has an equal probability of reaching the reactants and the products.^73^ Although this definition of a structural TS is not accurate, it is an illuminating way of extracting critical structures during the transition. In practice, we assume that the structural TS occurs when the *d*_rms_ values for the I and M phases are the same.^64^ *d*_rms_ is the root-mean-squared deviation of the pairwise distance (see the Supplementary Material for a definition).

There are two structural TS ensembles, one for each of the two directional transitions during the cell cycle. Overall, both TS ensembles have broad distributions. However, they are distinctly different from a structural perspective (Fig. 7). These findings are in contrast with those for the equilibrium process, since the TSs should be the same for the forward and backward reactions if the same condition is applied. For both TS ensembles, there was a slight decrease of PA1 relative to the I phase, unlike PA3 (Fig. 7(a)). The global structural order parameter *R*_*g*_ is similar for the two TS ensembles and has quite a narrow distribution. Moreover, Δ for the TS ensembles has a much broader distribution and a higher cylindrical-like value in the I → M than in the M → I transition (Fig. 7(b)). These findings indicate that the chromosomes in the two structural TS ensembles condense to a similar extent of compaction but have very different aspherical shapes.

**FIG. 7.**
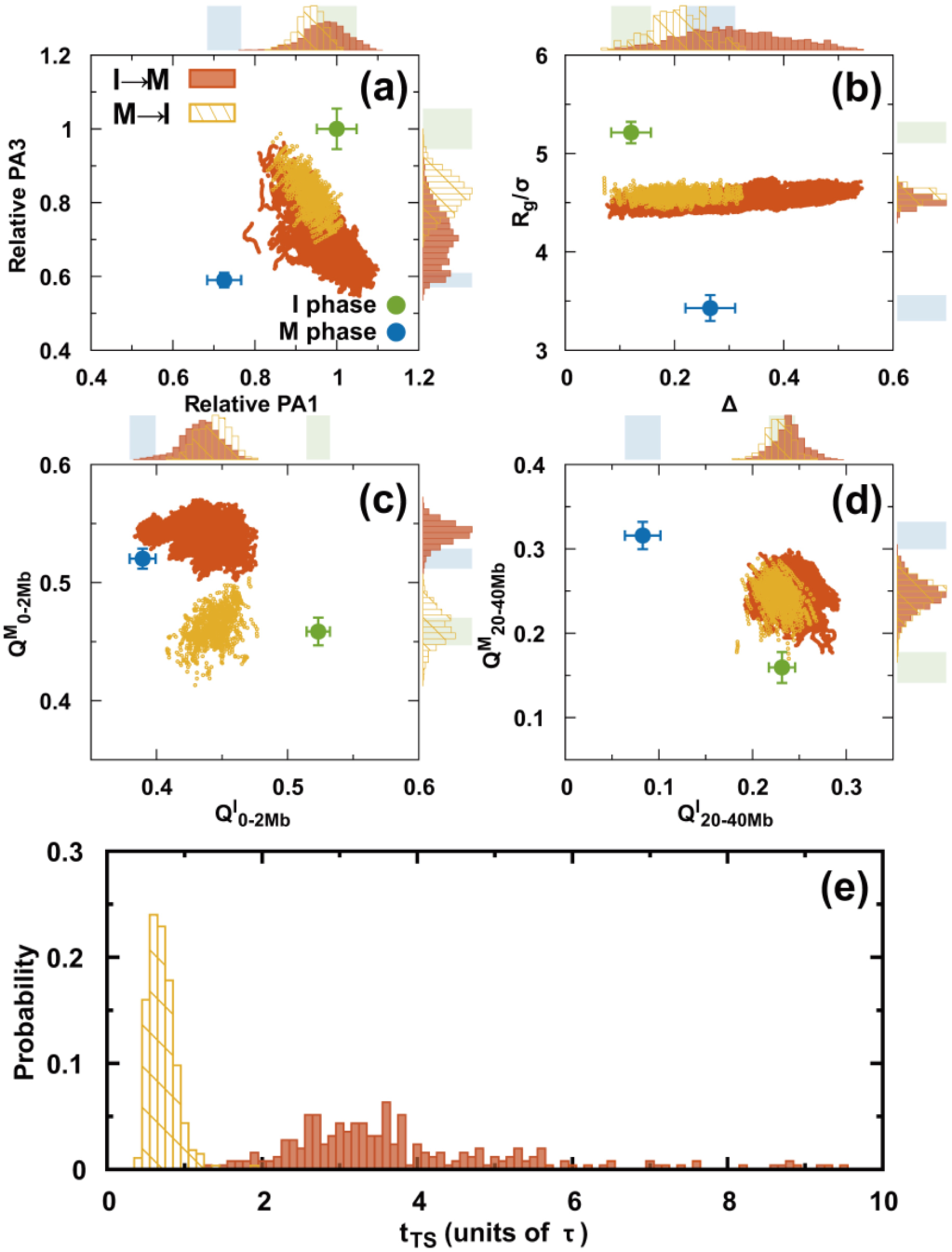
Structural TS ensembles. (a–d) The chromosomes in the TS ensembles are described by different order parameters. There are two TS ensembles for the transitions I → M and M → I. These are shown in orange and yellow, respectively. The plots with points in each panel show all the TSs in the trajectories. The top and right sidebars are the probability distributions projected onto different order parameters for one dimension. The shaded regions indicate the values for the I (green) and M (blue) phases. (e) Distributions of the first passage times to the TS ensembles.

From a contact perspective, the formation of the short-range interaction occurs closer to the final destined M phase than for the initial I phase in the structural TS ensembles of the I → M transition (Fig. 7(c)). However, an over-stabilized effect is observed at 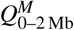. Moreover, the short-range contacts of the TS ensembles in the M → I transition are entirely distorted compared to those in the M phase and remain until nearly halfway through the transition to the I phase. In contrast, the long-range contacts in both ensembles resemble the formation in the I phase and are similarly located halfway to the M phase (Fig. 7(d)). Therefore, we deduced that the structural TS ensembles of the two directional transitions are mostly different from the interactions within the local TAD-length range and gradually become similar as the interaction range increases (Figs. S22 and S23).

There are also apparent differences between the two directional transitions in terms of the kinetic transition time (Fig. 7(e)). In the I → M transition, the first passage time to the structural TS ensembles (*t*_TS_) has a broader distribution along with a higher most probable value than in the M → I transition. We attribute the difference to the formation of the short-range contacts, as they contribute to the significant structural differences for the two structural TS ensembles.

### F. SMC complex-mediated loop formation during the cell cycle

The SMC complexes, such as cohesins and condensins, are critical in the reorganization of the chromosome structure during the cell cycle.^40^ In the loop extrusion model,^75^ these protein complexes are associated with the chromosome fiber and subsequently create a loop by progressively extruding. The process can lead to the accumulation of contact frequency in the Hi-C maps. The previous experimental studies used the slope of the logarithmic relation of contact probability *P*_*s*_ versus genomic distance *s* to infer the formation of cohesion and condensin loops.^26,74,76^ In practice, since *P*_*s*_(*s*) ≈ *s*^*α* (*s*)^, then *α*(*s*) was calculated as the slope of the logarithmic *P*_*s*_(*s*) versus *s*. A local maximum of *α*(*s*) in the local ranges (usually *<*2 Mb) indicates a deviation from the average decay, implying an enhanced contact probability for the SMC complexes. We calculated *s*(*α*) for the experimental Hi-C data in the I and M phases (Figs. 8(a) and 8(b)). The different positions of the local maximum in the *α*(*s*) curve indicate the sizes of the cohesin loop in the I phase and the condensin loop in the M phase, consistent with the previous analyses.^26,74,76^ Thus, we were able to delineate the loops formed from cohesin and condensin, though our model does not explicitly consider the protein–chromosome interaction.

**FIG. 8.**
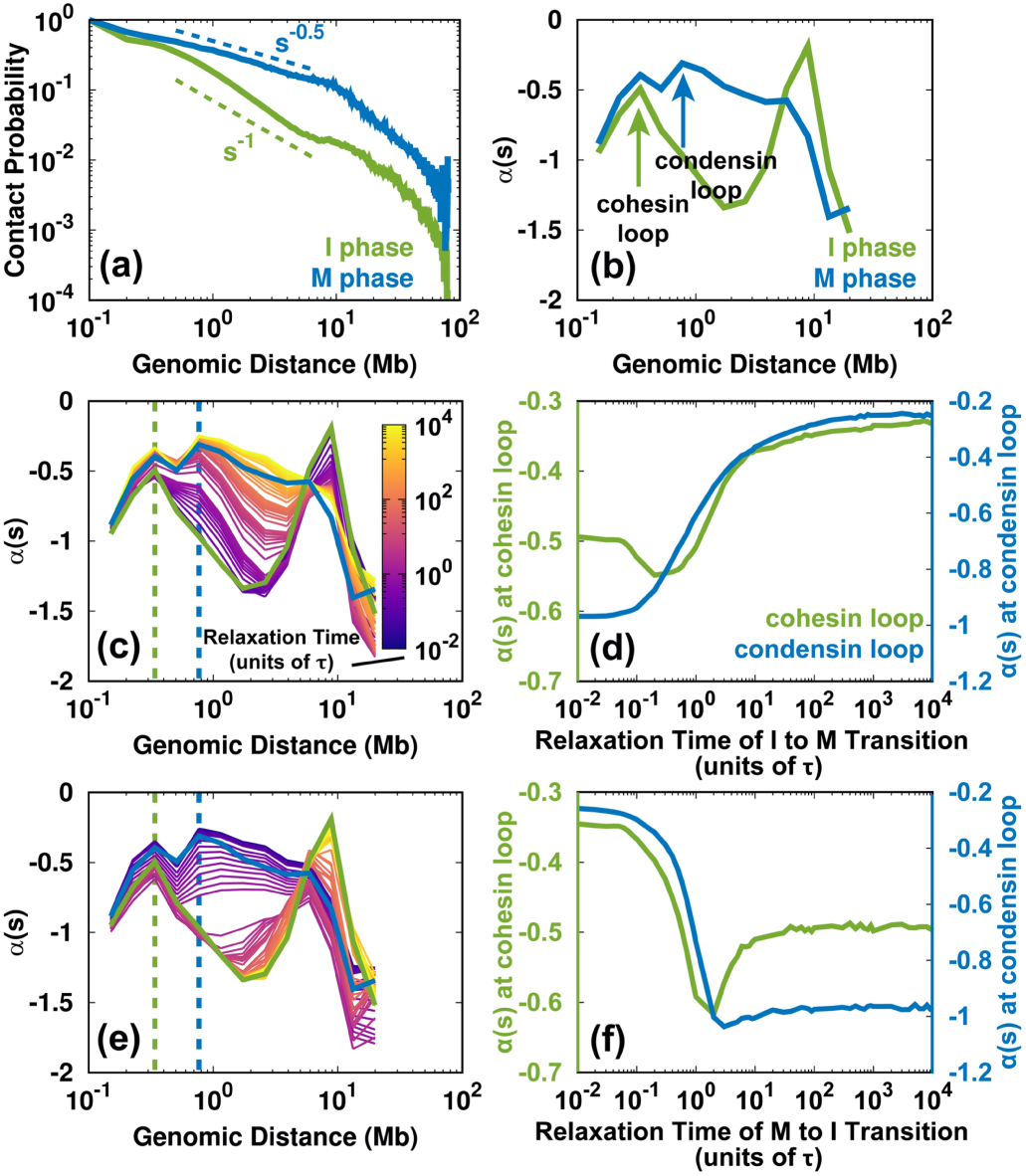
Changes to the SMC complex-mediated loops during the cell cycle. (a) Logarithmic plot of the contact probability *P*_*s*_ versus genomic distance *s* for the Hi-C data for the I and M phases. *P*_*s*_ has distinct decaying patterns in the I (∼−1) and M (∼−0.5) phases within 10 Mb. (b) The slope of *P*_*s*_(*s*) ≈ *s*^*α*(*s*)^ has local maxima in the I and M phases, corresponding to the sizes of the cohesion and condensin loops, respectively. There are two peaks in the M phase profile. The more significant maximum is at ∼800 kb, which corresponds to the loop formed by condensin.^26^ Condensin binding can also lead to enhanced contacts at short ranges (∼300 kb), similar to the range in which cohesin forms loops in the I phase.^26,74^ (c) Time evolution of *α*(*s*) for the I → M transition. The dashed lines indicate the sizes of the cohesin (green) and condensin (blue) loops. (d) The change of *α*(*s*) for the cohesin and condensin loops extracted from for the I → M transition. (e), (f) As (c) and (d) but for the M → I transition.

From the shape of the *α*(*s*) profile, we can identify a loss of the cohesin loop and a gain of the condensin loop in the I → M transition (Fig. 8(c)). Further projection of *α*(*s*) at the positions of the cohesin and condensin loops shows how these two complexes act during mitosis. As shown in Fig. 8(d), *α*(*s*) decreases at the position of the cohesin loop and increases at the position of the condensin loop at the very beginning of the I → M transition (up to 0.3*τ*). This is related to the simultaneous unbinding of cohesin and the binding of condensin. The late synchronous increase of *α*(*s*) for the cohesion and condensin loops is due to the mitotic compression during condensin binding, which increases the value of *α*(*s*) to that in the M phase.

On the other hand, the change of *α*(*s*) indicates the evolution of the formation and breakage of the cohesin and condensin loops for the M → I transition (Figs. 8(e) and 8(f)). Interestingly, the trends are not the reverse of those observed during the I → M transition. *α*(*s*) for both the cohesin and condensin loops decreases abruptly at the beginning of the transition until 2*τ*. This suggests the unbinding of condensin occurs without the binding of the cohesin. After 2*τ, α*(*s*) at the condensin loop has a relatively small change, implying that the condensin is entirely dissociated. In contrast, *α*(*s*) at the cohesin loop increases immediately after 2*τ*, indicating cohesin loading. In other words, our results imply that around 2*τ*, there is no association between cohesin and condensin in the chromosome. Notably, a recent experiment that focused on the mitotic exit process of HeLa cells detected a similar short-lived intermediate state, which was at the critical point for the cohesin-to-condensin (all-or-none) transition.^26^ We also found that at ∼2*τ* in the M → I transition, the chromosome has an over-expanded conformation. This is in agreement with our intuition that a chromosome in which cohesin and condensin do not bind can expand even more than in the I phase.

## IV. DISCUSSION AND CONCLUSIONS

We developed a landscape-switching model to investigate the chromosome conformational dynamics of the cell cycle. To capture the essence of the cell cycle, the model explicitly considers non-adiabatic non-equilibrium effects. In practice, the instantaneous energy excitation is used in the simulations, consistent with the evidence that the phase transition during the cell cycle occurs abruptly, like a switch.^50^ The rapid chromosome conformational relaxation in the post-switching landscape observed in simulations is consistent with experimental findings that the chromosome reorganization is fast during a phase transition.^19,26,27,68^ Importantly, we demonstrated the irreversibility of the chromosome conformational transition pathways during the cell cycle. Our findings confirm that the cell cycle is irreversible and controlled by different regulatory networks at different phases.^28^ Furthermore, we found that the significant irreversibility of the pathway is governed by the formation of local-range contacts when the transition is close to the I phase. By analyzing the formation of SMC complex-mediated loops, we provided a molecular-level explanation. The unloading of cohesin and the loading of condensin is simultaneous at the beginning of the I → M transition, in contrast to the pure loading of cohesin at the late stage of the M → I transition. Therefore, we suggest that the irreversibility of the chromosome conformational transition during the cell cycle can be attributed biologically to the different SMC complexes participating in the deformation (formation) of the I phase chromosome during the mitotic (exit) process.

The description of chromosome mitotic folding obtained from the simulated pathways is consistent with the previous experimentally proposed two-stage hypothesis.^19,21^ From Fig. 3(a), we can estimate when the first step of linear compaction ends and the second step of axial compression begins as when the slope of the average pathway on the PA extension reaches the half-value (∼20*τ*). The condensin loop is 90% formed at ∼20*τ* in our simulations (Fig. 8(d)). This description of concurrent linear compaction and the formation of a condensin loop is consistent with the experimental observation that linear compaction is led by the condensin-mediated consecutive chromosomal loop.^19,21^ At the second stage, the chromosome undergoes axial compression associated with an increase in the contact probability at a longer range. Based on our kinetic analysis (Fig. 6(c)), the anisotropic chromosome condensation from the I to the M phase is likely an outcome of the asynchronous formation of contacts at various genomic distances, as the local contacts form much faster than the non-local ones. In addition, we produced a description of the mitotic exit process that is consistent between the simulations and experiments.^26^ We found that the mitotic exit was efficient for the over-expanded chromosome conformation at ∼2*τ*, when the chromosome is also free of cohesin and condensin binding. The transient state at ∼2*τ* in our simulations captures the main characteristics of the experimentally uncovered intermediate state that dominates the condensin-to-cohesin transition.^26^ Topologically, the over-expanded chromosome has more surface accessible to the SMC complexes, which biologically enhances the subsequent cohesin targeting. Therefore, our structural-level description suggests that there is a biological role for the aberrant intermediate state.^77^

Our investigations of the cyclic structural changes of the intra- and inter-TADs provide detailed descriptions of TAD condensation and insulation during the cell cycle. Consistent with the recent single-cell Hi-C experiments,^18^ the decondensed TADs were largely observed in the I phase, suggesting that open chromatin is functionally advantageous to gene regulation.^12^ Additionally, it was experimentally observed that some TADs can de-condense, although the overall trend for TAD condensation has a monotonic increase. In our simulations, we observed such deviations for particular TADs and we confirmed the heterogeneity in the TAD transformations. On the other hand, the insulation of a TAD evolves more cooperatively than TAD (de-)condensation does, implying that the spatial proximity of neighboring TADs is an indicator of cell-cycle progression.^18^

It is tempting to match our kinetic simulations to the real cell-cycle process based on the current experimental findings. For mitosis, experiments found that TADs disappear in the prophase. By using the decrease of the TAD signal variance as an indicator for the loss of the TAD boundary, we deduced that the prophase occurred at ∼20*τ*, when 90% of the TADs are lost (Fig. S11). This is coincidentally the critical point between the two stages of mitotic folding. This time point can be used to separate the first stage of condensin-dependent linear looping in the prophase^78^ from the second stage of axial compression in the prometaphase.^79^ In our simulations, the further relaxation of the chromosome from the prometaphase to the metaphase can take up to 1000*τ* or more, which is consistent with the experimental findings that the prometaphase is long.^21^ On the other hand, experiments detected the aberrant intermediate state in the telophase during the mitotic exit process. We determined that the telophase starts at ∼2*τ*. Moreover, we observed the establishment of TADs at 2*τ* in our simulations (Fig. S11), in agreement with the experimental analysis that TADs initially form as early as the telophase. In addition, a relatively small change in the simulation trajectories can be seen after 10*τ* (Fig. S17), indicating the time of the entry into the I phase. Our observation of compartmentalization proceeding after 10*τ* (Fig. S13) resonates with the experiments, which showed that the strengthening of compartments can continue for a long time in the I phase.

Overall, our simulations have predicted well the key progression phases in chromosome morphogenesis during the cell cycle. However, there are two things worth noting:

1. The underlying kinetics of the system is based on landscape switching. The timescale estimated in our simulations may not be directly linked to the real reaction rates.
2. The additional axial compression of the chromosome in the anaphase after the metaphase^80^ was not observed in our model. Making predictions for the anaphase is beyond the scope of this simulation, as we have roughly assumed that the metaphase chromosome was the final destined condensed state.

Energy landscape theory has successfully been used to investigate the folding and function of proteins.^81,82^ The extension of the energy landscape concept to describe genomic dynamics seems sensible. However, it depends on the application.^83^ Beyond the thermodynamic distributions, our study and other previous studies showed that effective energy landscapes can correctly characterize the kinetics in chromosome anomalous diffusion.^67,84^ These features suggest the broad applicability of the effective energy landscape approach for much bigger and more complex systems beyond proteins, i.e., chromosomes. Nevertheless, there are questions. For example, can the effective energy landscape properly describe the chromosome conformational transition during the cell cycle? A direct application of the energy landscape seems to be inappropriate, as the driving forces in the cell cycle originate mainly from non-equilibrium chemical reactions, such as ATP hydrolysis. However, the transition process can be approximately represented as a switch between the two phases, such that the prior and posterior energy landscapes in each phase are still effective and correct for describing the chromosome dynamics within the phase. The performance of the energy landscape-switching model between cell-cycle phases was assessed by comparing the mitotic and mitotic exit chromosome folding results from experiments and our simulations. The emergent descriptions from our simulations are very consistent with those observed in experiments, supporting the validity of our model. The irreversibility of the pathways uncovered in our simulations is an outcome of the non-equilibrium dynamics,^85^ and would be absent if equilibrium dynamics were applied. Therefore, our work suggests that combining energy landscapes with non-equilibrium effects is effective at the genomic level.

In summary, we have developed quite a complete physical description of cell-cycle chromosome dynamics. Our results may aid the global and quantitative understanding of functional chromosome reorganization in cell development and the pathological disorganization of cancer cells. Our detailed work on the roles of SMC proteins elucidates the genome structure and dynamics at the molecular level. Thus, it may provide useful guidance in practical research on modulating and controlling the cell cycle, which is key to understanding cancer and cellular senescence. We anticipate that our model could lead to new approaches for developing molecular simulations of higher-level biological processes, which are mostly non-equilibrium.

## SUPPLEMENTARY MATERIAL

See the Supplementary Material for definitions of the order parameters, the methods for identifying the TADs and calculating the compartment profile, and additional figures.

## ACKNOWLEDGMENTS

We thank the National Science Foundation (Grant Nos. PHY-76066 and CHE-1808474) and the National Institute of General Medical Sciences of the National Institutes of Health (Grant No. R01GM124177) for their support. The content is solely the responsibility of the authors and does not necessarily represent the official views of the National Institutes of Health.

## DATA AVAILABILITY STATEMENT

The data that support the findings of this study are available from the corresponding author upon reasonable request.

