## Supplementary Material for "Conformational state switching and pathways of chromosome dynamics in cell cycle"

### Supplementary Material
for
Conformational state switching and pathways of chromosome dynamics in cell cycle

Xiakun Chu [1], and Jin Wang[1,2*]

[1] Department of Chemistry
[2] Department of Physics and Astronomy
State University of New York at Stony Brook, Stony Brook, NY 11794, USA

*

### Quantities and order parameters

We use the extensions of principal axes (PA), $\Delta$ and $S$ suggested previously to determine the shape of the chromosome polymer chain [1, 2]. These quantities are determined by the inertia tensor $\boldsymbol{IT}$, which can be written as follows:

$$\boldsymbol{IT} = \begin{bmatrix} \boldsymbol{r_x r_x^T} & \boldsymbol{r_x r_y^T} & \boldsymbol{r_x r_z^T} \\ \boldsymbol{r_y r_x^T} & \boldsymbol{r_y r_y^T} & \boldsymbol{r_y r_z^T} \\ \boldsymbol{r_z r_x^T} & \boldsymbol{r_z r_y^T} & \boldsymbol{r_z r_z^T} \end{bmatrix}$$

, where $\boldsymbol{r_x}$, $\boldsymbol{r_y}$ and $\boldsymbol{r_z}$ are the row vectors for the positions of genomic loci shifted by the corresponding means. The eigenvalues ($\lambda_k$) of $\boldsymbol{IT}$ correspond to the squares of the extension lengths along the three PA. Thus, $R_g^2 = tr\boldsymbol{IT} = \sum_{k=1}^{3} \lambda_k$. Then we have aspherical quantity $\Delta$:

$$\Delta = \frac{3}{2} \frac{\sum_{k=1}^{3} (\lambda_k - <\lambda>)^2}{(tr\boldsymbol{IT})^2}$$

, and $S$:

$$S = 27 \frac{\prod_{k=1}^{3} (\lambda_k - <\lambda>)}{(tr\boldsymbol{IT})^3}$$

Deviation of $\Delta$ from 0 indicates the extent of anisotropy. Negative and positive values of $S$ correspond to oblate and prolate shapes, respectively.

We use the fraction of contact $Q$, which was previously used in protein folding simulation [3], to monitor the structural similarity of two chromosomes. By using the average pairwise distances in the chromosome at the I or M phase shown in Fig. S4 and S5, we can compare one chromosome structure to the ensemble of either I or M phase. In practice, $Q$ is defined as:

$$Q = \frac{1}{N_{sum}} \sum_{i,j} exp(-\frac{(r_{i,j} - < r_{i,j}^{I(M)} >)^2}{2\delta^2})$$

, where $N_{sum}$ is the number of summed pairs, $< r_{i,j}^{I(M)} >$ is the average distance between genomic locus $i$ and $j$ of the I (M) phase ensemble, and $\delta$ controls the slope of the function with the value of $\delta = 0.5\sigma$.

To see how contacts change with different genomic distances, we calculate the average $Q_l^{I(M)}$ and contact probability $P_l$ within the defined genomic distance separation $l$. Therefore, $Q_l^{I(M)}$ and $P_l$ are expressed as:

$$Q_l^{I(M)} = \frac{1}{N_{sum}} \sum_{i,j}^{|i-j| \in l} Q_{i,j}^{I(M)}$$

$$P_l = \frac{1}{N_{sum}} \sum_{i,j}^{|i-j| \in l} P_{i,j}$$

, where a variety of $l$ is practically applied: 0-2Mb, 2-5Mb, 5-10Mb, 10-15Mb, 15-20Mb, 20-40Mb and total (0-81.2Mb).

We further use the root-mean-square deviation of the pairwise distance ($d_{rms}$) to quantify the structural differences between one chromosome and the ensemble of the I or M phase. $d_{rms}^{I(M)}$ is expressed by:

$$d_{rms}^{I(M)} = \sqrt{\frac{1}{N_{sum}} \sum_{i,j} (r_{i,j} - < r_{i,j}^{I(M)} >)^2}$$

We use $d_{rms}$ between two chromosome structures to perform clustering at the I and M phase after the maximum entropy principle simulations. Practically, we use a hierarchical clustering algorithm suggested in [4]. The algorithm works as follows. Firstly, we set up a cut-off value $d_{rms}^c$, if the $d_{rms}$ between two chromosome structures is smaller than the $d_{rms}^c$, we consider these two structures similar and group them into the same cluster. Then we increase the value of $d_{rms}^c$ and repeating the grouping process. Finally, we have a hierarchical clustering dendrogram for the I phase chromosome ensemble shown in Fig. S8 and the M phase chromosome ensemble shown in Fig. S9. We find that when $d_{rms}$ reaches $3.0\sigma$ at the I phase and $2.0\sigma$ at the M phase, the contact maps of the chromosome structures between different clusters are distinguishable (Fig. S8 and S9), but show high similarity within the same cluster. This indicates a good criterion to determine the $d_{rms}^c$. Further kinetic simulations show the chromosome dynamics within both the I and M phase is very slow, and transition between clusters cannot be achieved (Fig. S6 and S7). This feature demonstrates the validity of using the representative chromosome structures from clusters as the initial conditions to perform the phase-transition simulations using the energy landscape-switching model.

### TAD and compartment identifications

The TAD signals from the simulations are calculated by log2 ratio of the contact probabilities from upstream-to-downstream within 2Mb regions, same with the previously used methods [5, 6]. We additionally use the insulation score, suggested by Crane et al. [7], to describe the formation of the TAD boundary. We use the same size of sliding square (500×500kb) within the original application to calculate the insulation score [7].

The calculation of compartment profile follows the method proposed in [8], with minor modifications. The compartment profile is usually from one of the top-weighted components of the contact probability matrix and can be calculated via principal component analysis (PCA) [9]. We first calculate the observed/expected contact probability matrix at 1Mb resolution. The expected matrix is calculated as the mean contact probability at a given genomic distance, same as suggested by Nagano et al. [10]. We then convert the $P_{obs}/P_{exp}$ to a Pearson correlation matrix, which is subsequently used for PCA ($P_{obs}$ and $P_{exp}$ are observed and expected contact probability at 1Mb resolution, respectively.). For the long segments of chromosome 5 we used here, we find that the first PC (PC1) corresponds to the plaid pattern and thereafter is used for the compartment profile. The direction of the PC1 values is arbitrary, and we set the positive and negative PC1 values with gene density (positive to gene-rich and negative to gene-poor). This is done at the interphase Hi-C data. The direction of the PC1 values during the cell cycle is determined according to the correlation coefficient with the PC1 values of the interphase Hi-C data.

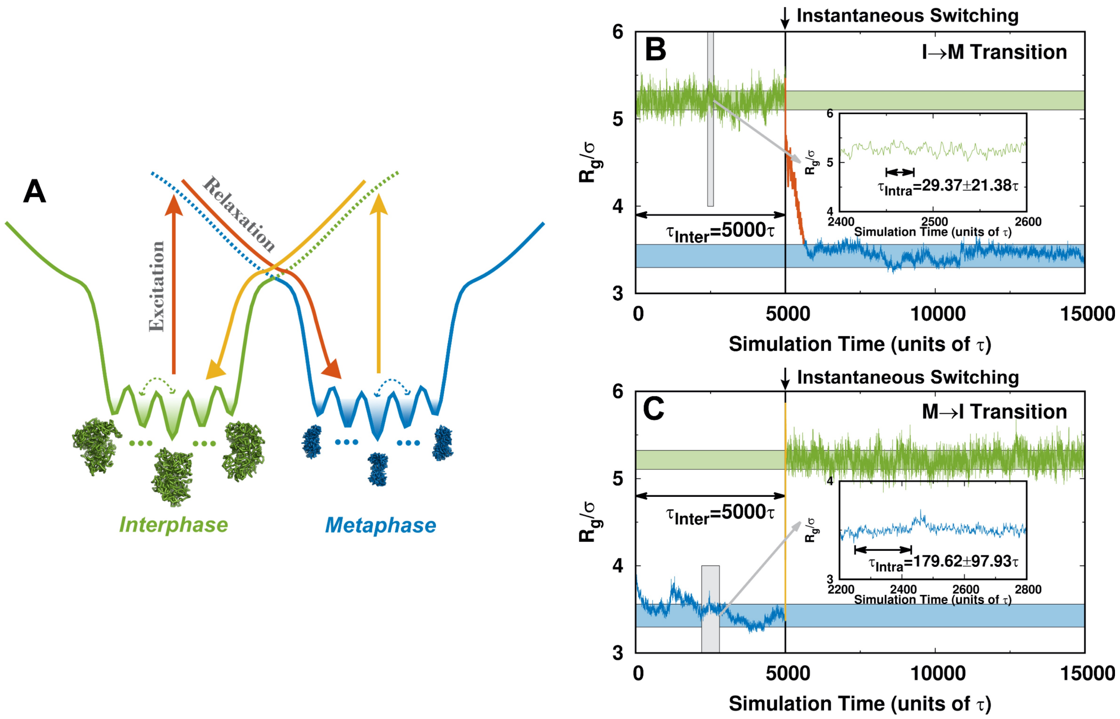


**FIG. S1:** The non-adiabatic non-equilibrium landscape-switching model and simulations. (A) The illustration of the landscape-switching model, the same as Fig. 1 in the main text. (B) One representative trajectory of the I→M transition. The radius of gyration $R_g$ is used as the order parameter for monitoring the chromosome conformational dynamics. The simulation starts from one chromosome structure at the I phase. The time for simulating the chromosome at the I phase is set to $5 \times 10^3 \tau$, after which the instantaneous switching from the I to M phase is implemented. The instantaneous switching leads to an infinite rate for the phase-transition, behaving as a switch. The waiting time for dwelling at the I phase is $5 \times 10^3 \tau$, which corresponds to the inter-landscape hopping time. The time for the intra-landscape dynamics is estimated by the decay of the autocorrelation function of $R_g$ [11]. We observe that the timescale of the inter-landscape hopping is much slower than that of the intra-landscape dynamics. Besides, only one switching is performed in each trajectory in order to accord with the fact that cells spend significant time dwelling at one phase during the cell cycle process, so multiple energy excitations are unrealistic. Therefore, the simulation leads to an extreme of non-adiabatic process. The insert plot is the zoom-in trajectory at the I phase. (C) The same as (B) but for the M→I transition.

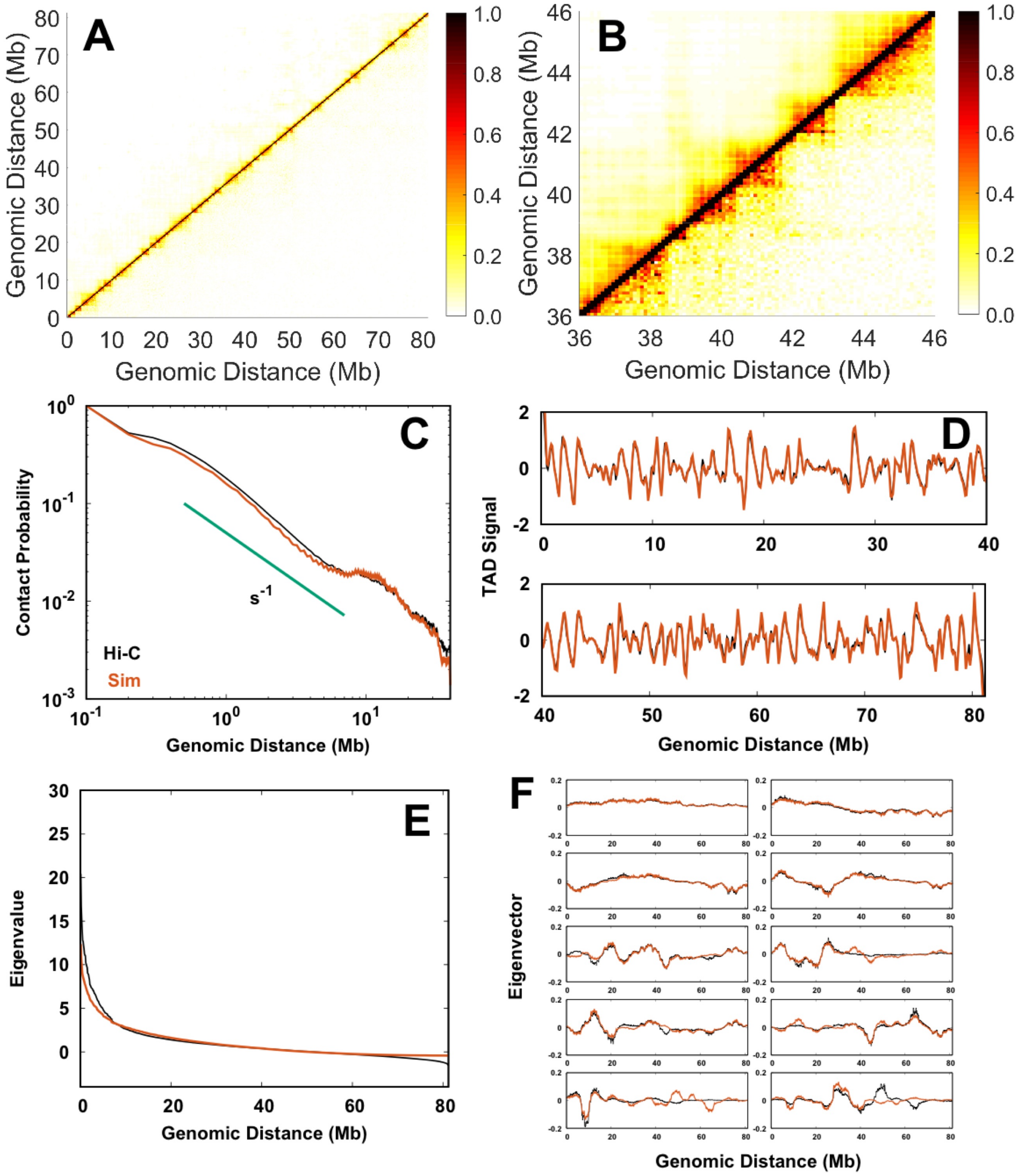


**FIG. S2:** Comparison of the chromosome ensemble at the I phase between simulation and Hi-C experiment. (A) The 2D probability heat map. The top left is the simulation data, and the bottom right is the Hi-C data. The correlation coefficient between the full simulation and experiment datasets is 0.91. (B) Zoom-in 2D contact probability heat map. (C) The contact probability along with the genomic sequence separation distance. (D) The TAD signal. (E) The eigenvalues of the contact probability matrices. (F) The top 10 eigenvectors of the contact probability matrices. In (C-F), the orange and black lines represent the quantities calculated from the simulation and Hi-C data, respectively.

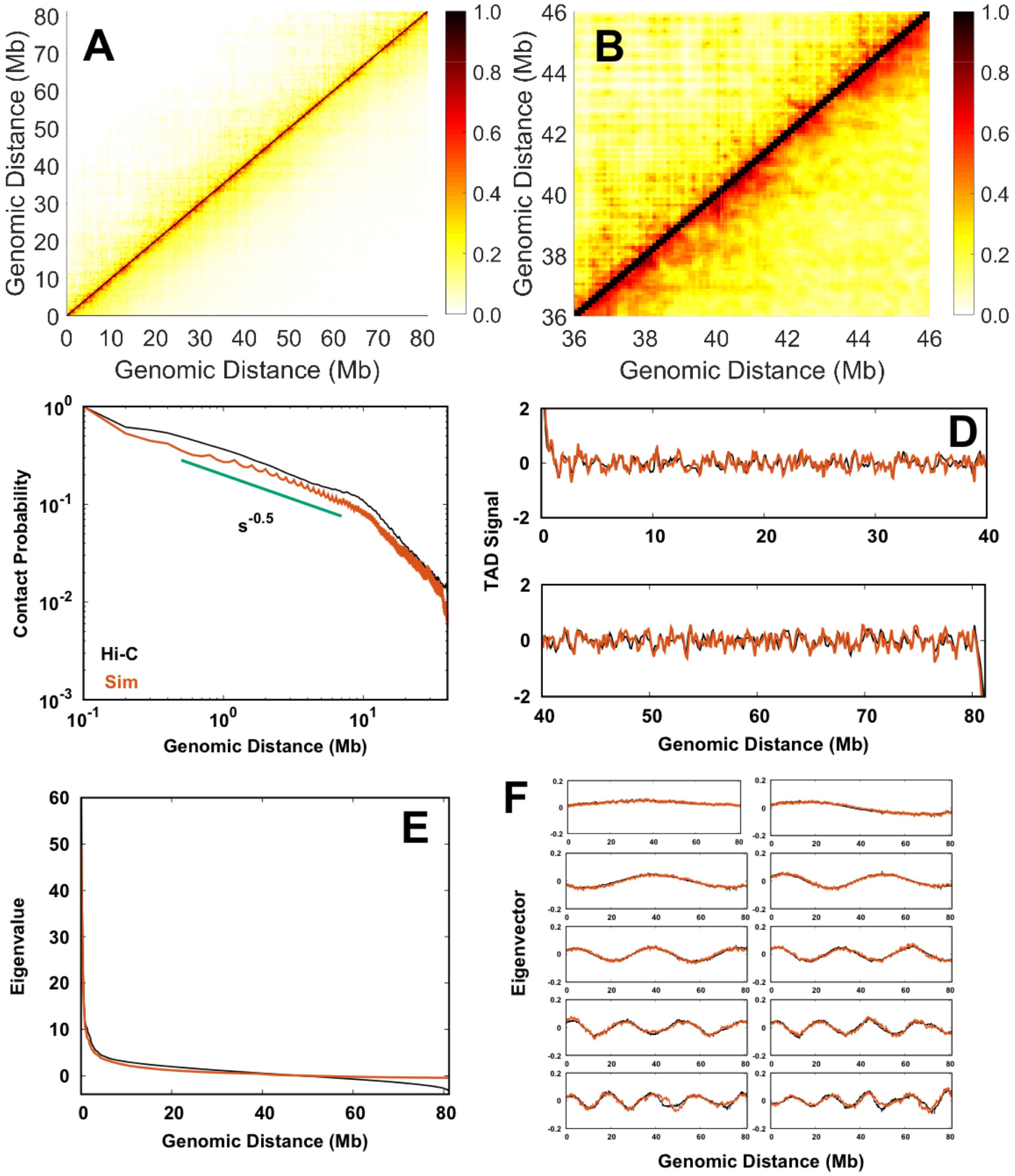


**FIG. S3:** Comparison of the chromosome ensemble at the M phase between simulation and Hi-C experiment. Plots are the same with those in Fig. S2. In (A), the correlation coefficient between the full simulation and experiment datasets is 0.91.

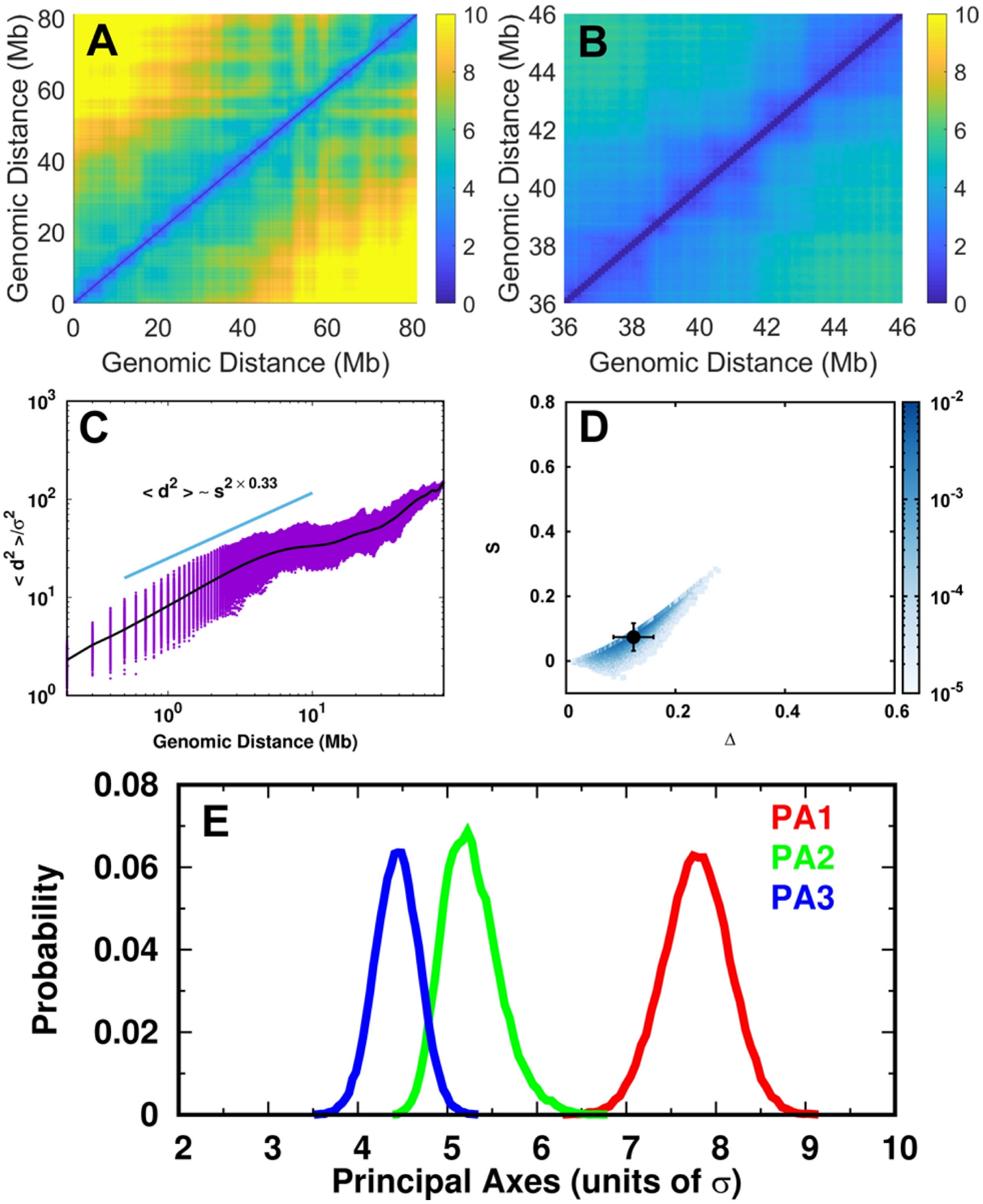


**FIG. S4:** Physical spatially structural organization of chromosome at the I phase. (A) The 2D locus-locus distance $d$ map. $d$ is in the reduced length unit of $\sigma$. (B) Zoom-in 2D locus-locus distance $d$ map. (C) The second-order movements of the locus-locus distance $d^2$ along with the genomic sequence separation distance. (D) The aspheric shape parameters. The black point with error bars presents the mean value with its variances. (E) The distributions of the extension lengths along the three principal axes.

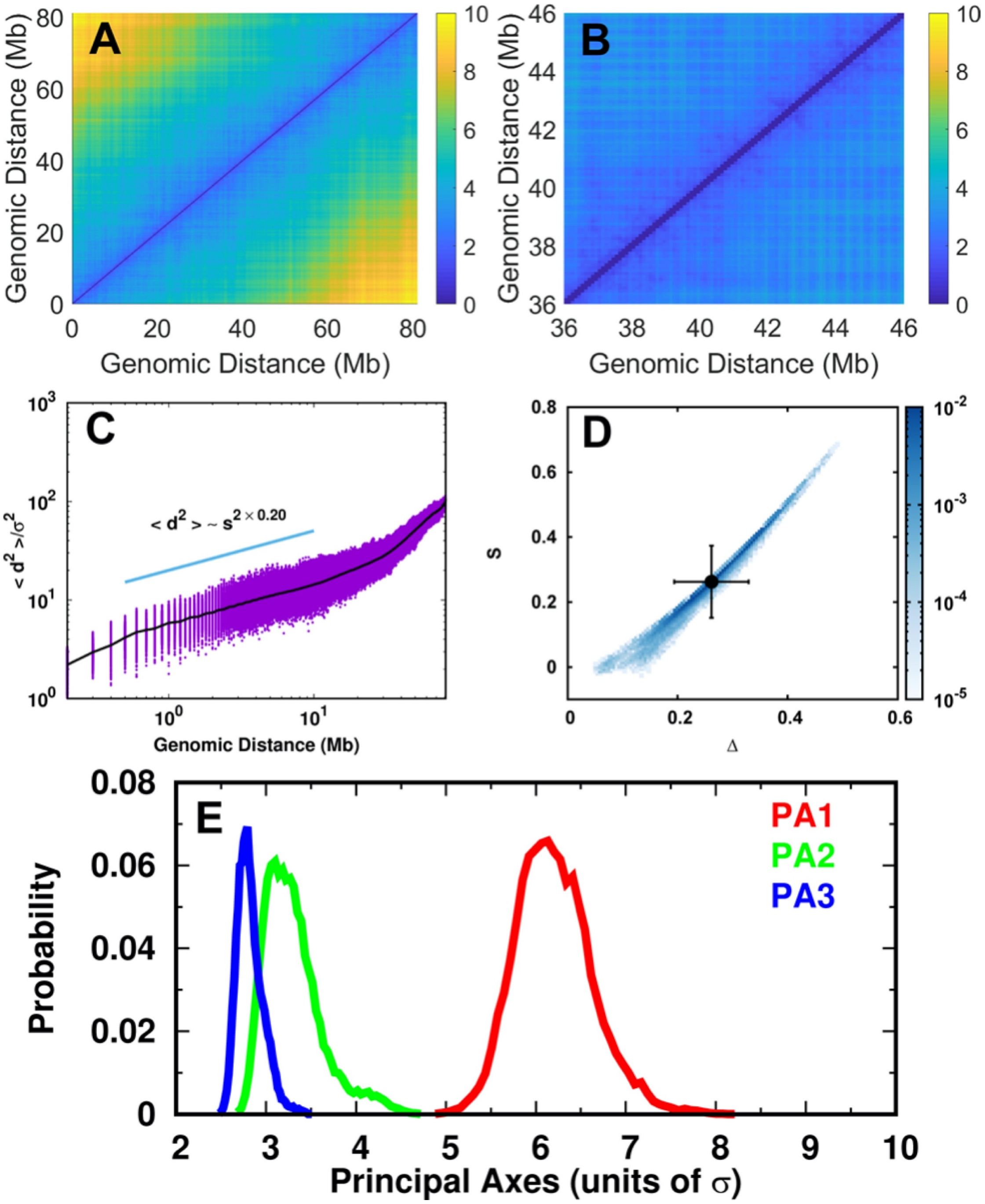


**FIG. S5:** Physical spatially structural organization of chromosome at the M phase. Plots are the same with those in Fig. S4.

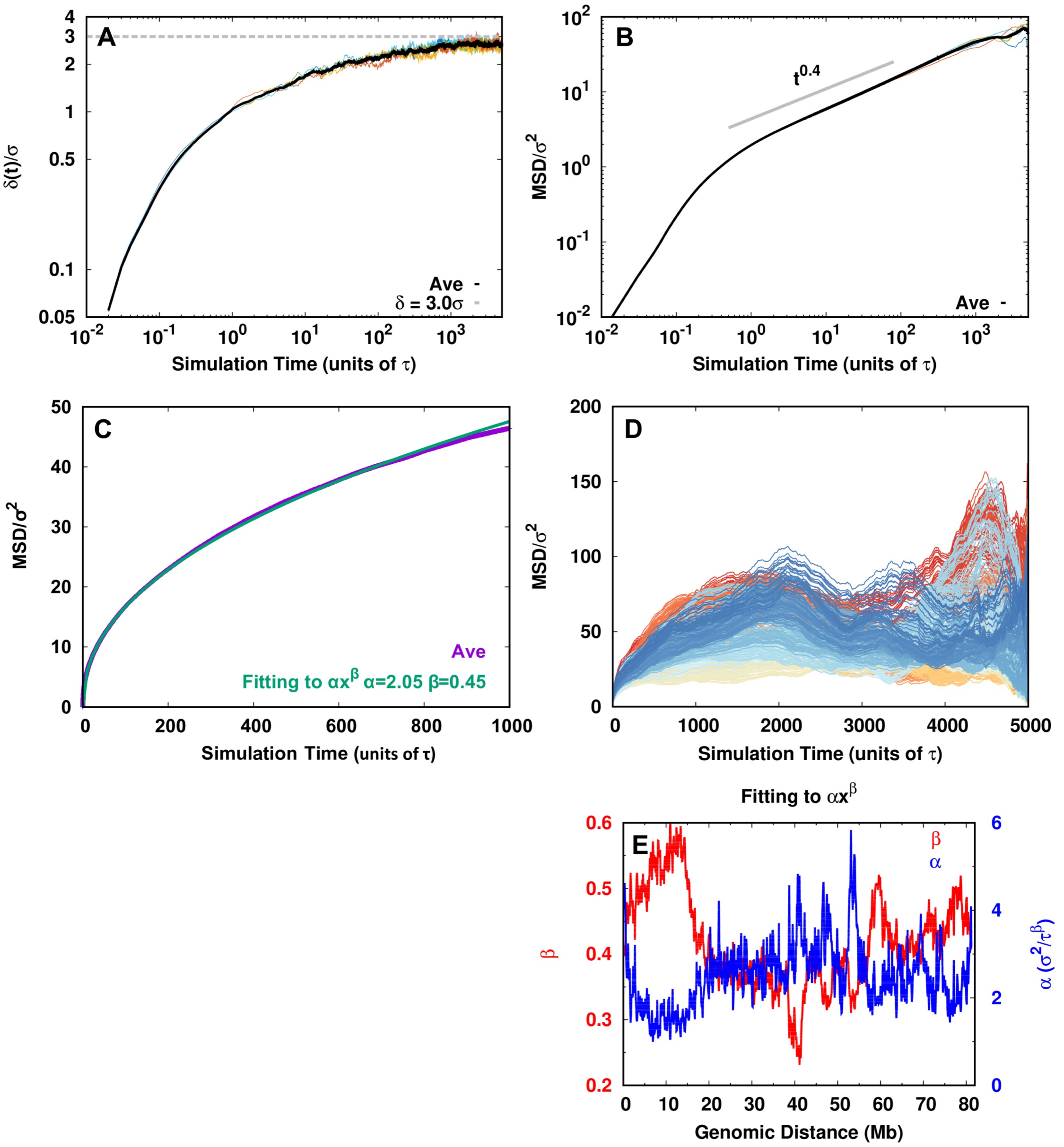


**FIG. S6:** Chromosome conformational dynamics at the I phase. (A) The time evolution of the average $d_{rms}$ between every loci pair at the time $t$ relative to its initial value $\delta(t) = \sum_{i,j} d_{rms}(i,j,t)/N_{pairs}$, where $N_{pairs}$ is the number of the summed pairs. (B) The average mean square displacement (MSD) for all the genomic loci. (C) Fitting of MSD to the power-law function. (D) MSD of the individual genomic locus. (E) The fitting parameters of MSD for individual genomic locus. Data in (A-C) are extracted from five trajectories, each starting from the center structure of the top five population-occupied clusters. Data in (D, E) are from the trajectory starting from the center structure of the largest cluster. The chromosome at the I phase exhibits sub-diffusive characteristics and high heterogeneity for different individual loci. To estimate the physical unit of our model, we can approximately regard our 100-kb coarse-grained bead as 30-nm fiber, which directly gives the Kuhn length unit $\sigma \approx 150nm$. The experimental MSD of chromosome in HeLa cell has been measured by the single nucleosome imaging technique that shows MSD at $0.5s$ is in the range of 0.01-0.015$\mu m^2$ [12]. Mapping our simulation data to experimental measurement leads to $\tau \approx 2s$. Therefore, our longest simulation can reach as long as $\tau_{max} = 1 \times 10^4 \tau \approx 5.5$h in the real-time.

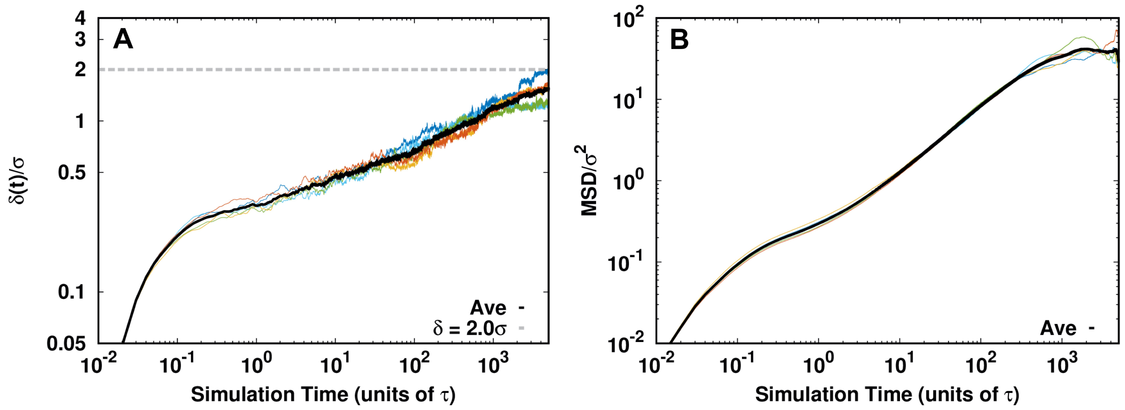


**FIG. S7:** Chromosome conformational dynamics at the M phase. (A) The time evolution of $d_{rms}$ between every locus pair at the time $t$ relative to its initial value $\delta(t)$. (B) The average MSD for all genomic loci.

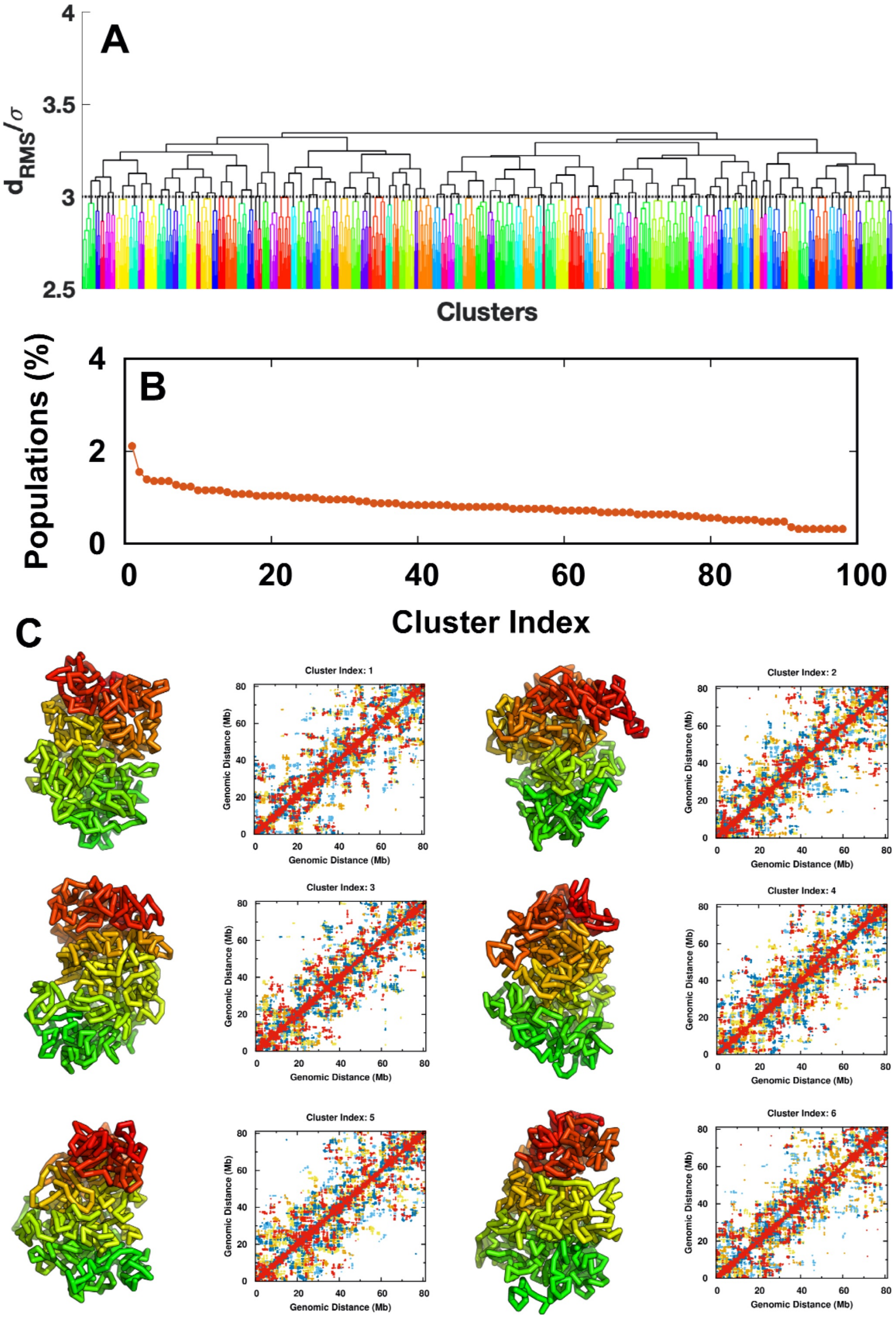


**FIG. S8:** Chromosome clustering and structural illustrations at the I phase. (A) The hierarchical clustering shown as a dendrogram. Cut-off distance 3.0$\sigma$ is applied. (B) The cluster populations. (C) The top 6 population-occupied clusters. Each structure is picked from the center of the corresponding cluster. The contact map is shown by a mixture of 5 structures within the cluster.

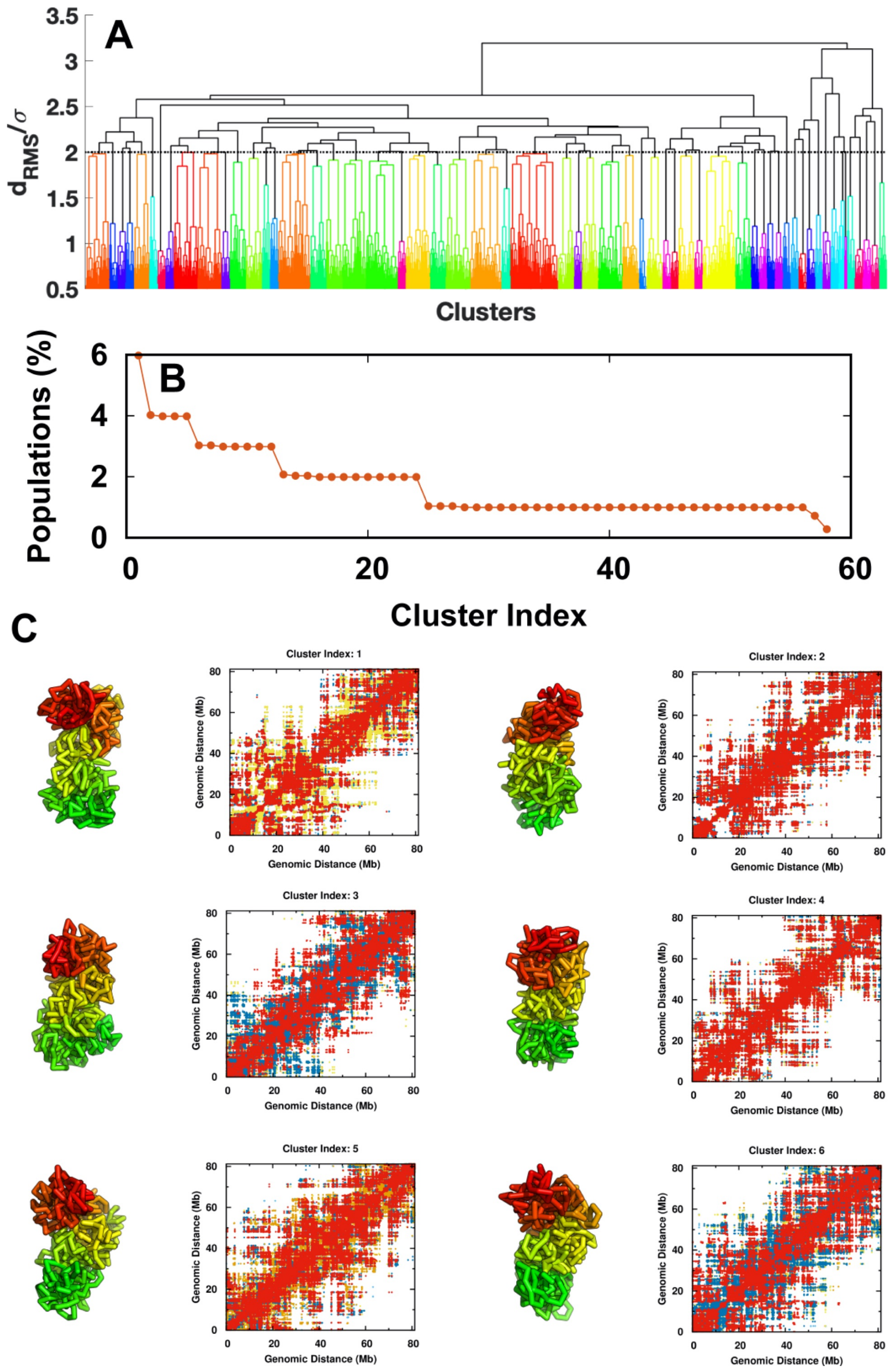


**FIG. S9:** Chromosome clustering and structural illustrations at the M phase. Plots are the same with those in Fig. S8. The cut-off distance is $2.0\sigma$.

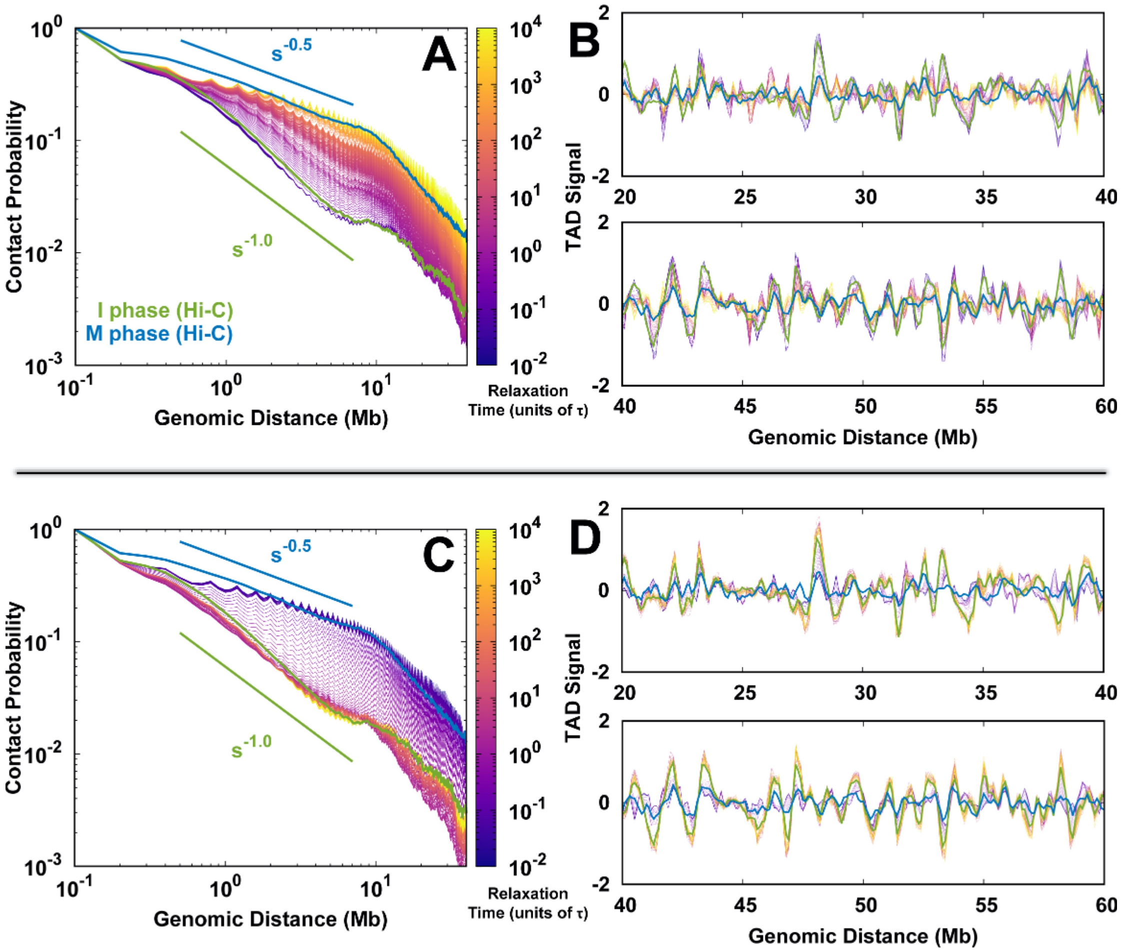


**FIG. S10:** Chromosome conformational transition during the cell cycle. The chromosome conformational evolution from the I to M phase is shown in terms of (A) the contact probability and (B) the TAD signal along with the genomic distance. The color schemes of lines dictate the time proceeding in logarithmic scale. The green and blue lines are the corresponding quantities directly obtained from the experimental I and M Hi-C data, respectively. (C, D) The same as (A, B) but for the transition from the M to I phase.

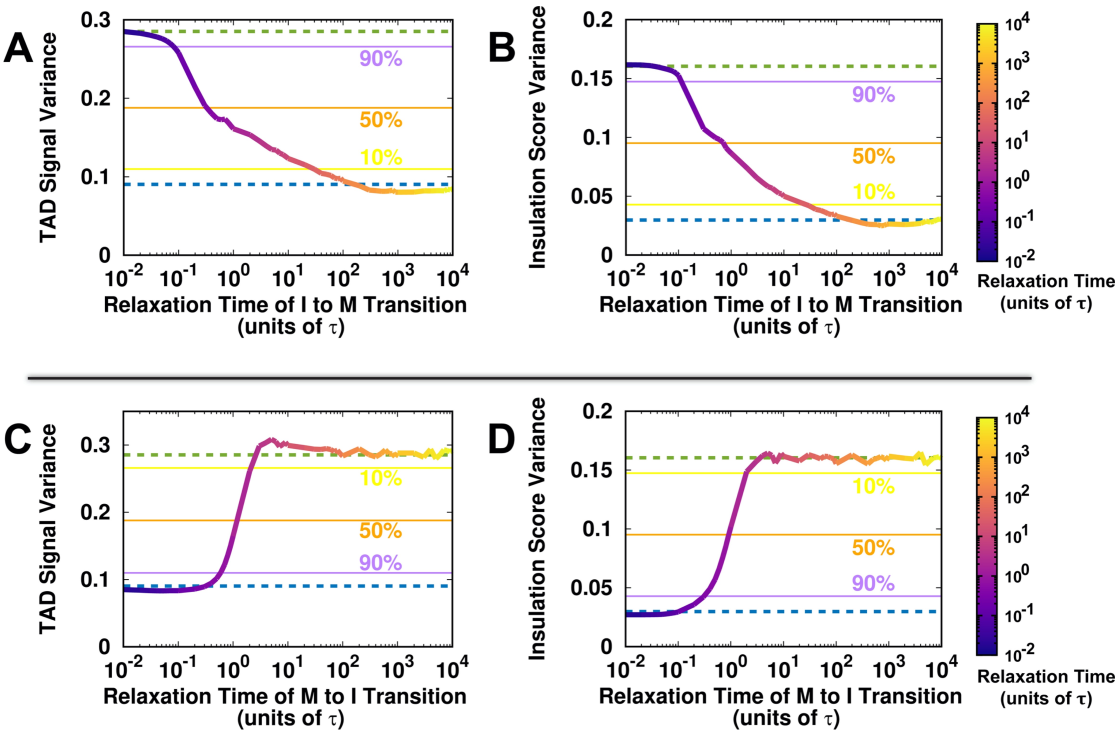


**FIG. S11:** The evolutions of (A) the TAD signal variance and (B) the insulation score variance of the chromosome during the I to M transition. Insulation score indicates the amount of contacts formed across a chromosomal locus to a certain distance. TAD boundaries have a low score (indicative of high insulation), whereas loci inside TADs show a high score (little insulation). Both the TAD signal variance and insulation score variance provide a quantitative measure of the presence of TADs (the large variance corresponding to the high propensity of TAD formation). The purple, orange and yellow lines are used as indicators for the initial loss of TADs (90% of variance conserved), half loss of TADs and entire loss of TADs (10% of variance conserved), respectively. The green and blue dashed lines indicate the values at the I and M phase, respectively. (C, D) The same as (A, B) but for the transition from the M to I phase.

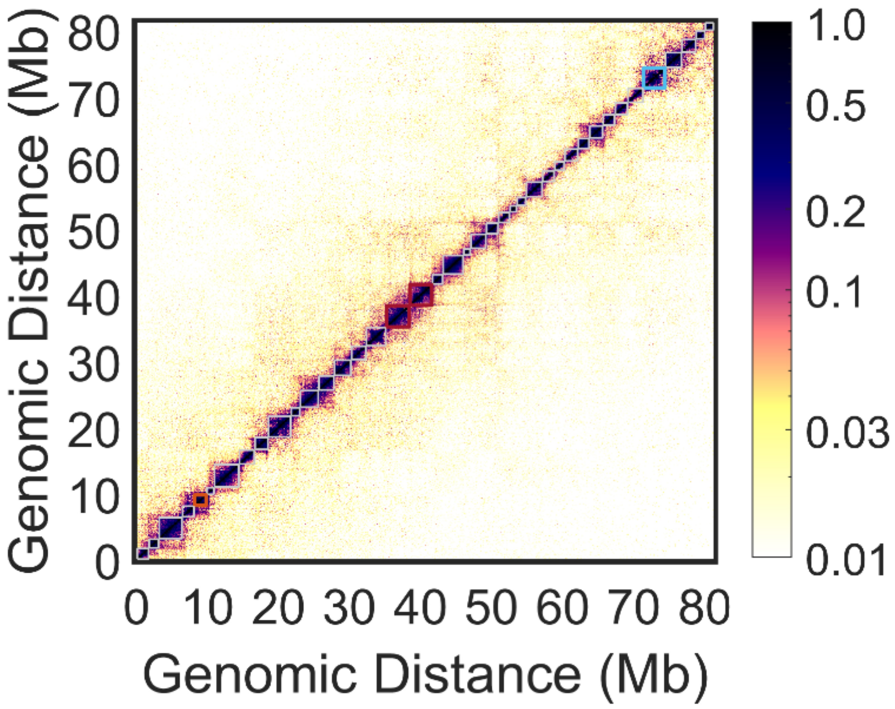


**FIG. S12:** TADs identified by the HiCSeg software [13]. There are 40 TADs detected at the I phase Hi-C data with an average size of 1.93 Mb and marked as grey squares on the Hi-C heat map. TAD5 (8.2-9.8Mb) and TAD36 (71.4-74.4Mb) are colored orange and blue respectively, and discussed in the main text for the intra-TAD structural evolution during the cell cycle. TAD17 (35.2-38.4Mb) and TAD18 (38.5-41.6Mb) are colored red and discussed in the main text for the inter-TAD structural evolution during the cell cycle.

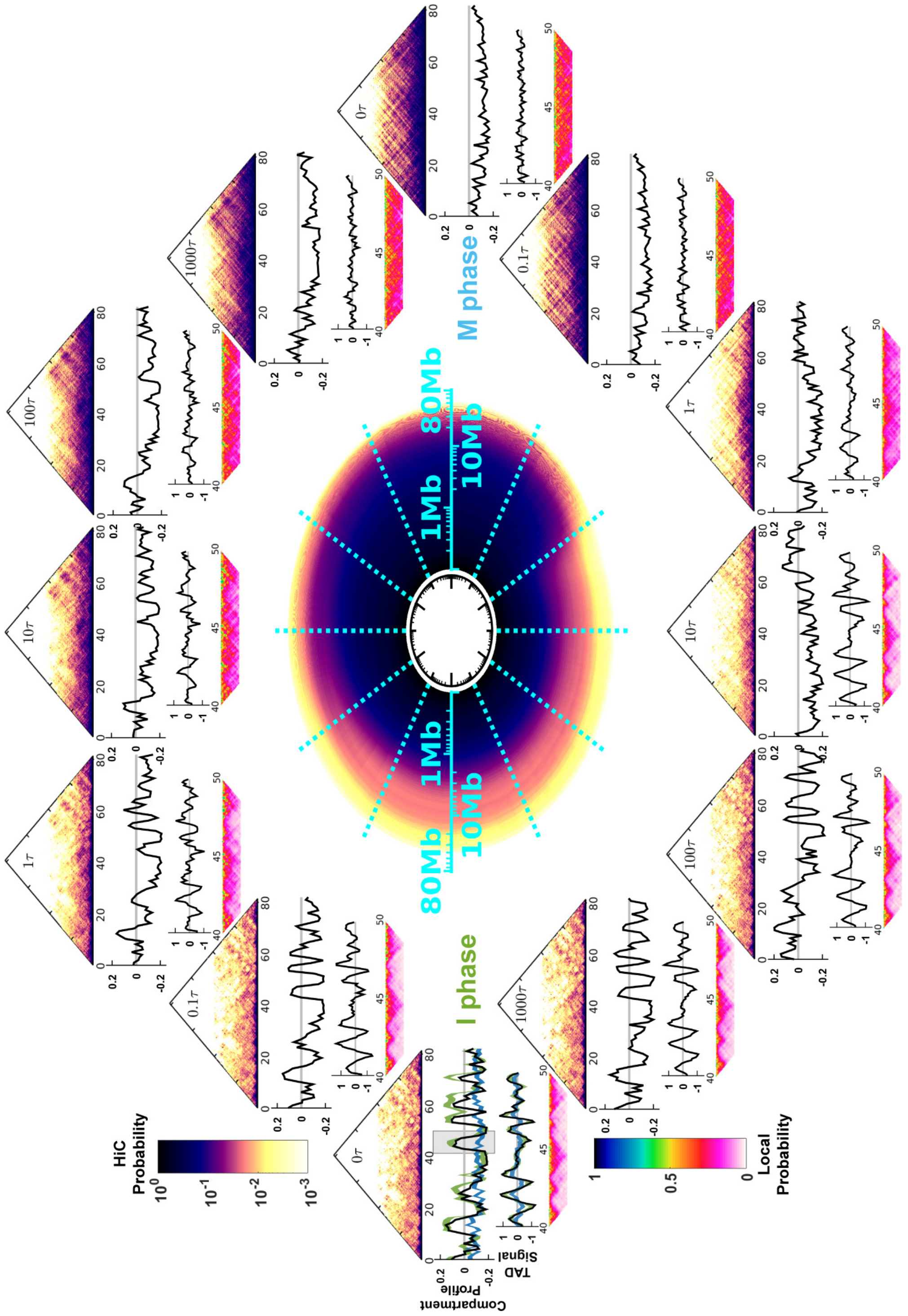


**FIG. S13:** The same with Fig. 2 in the main text with three addition time points ($t = 0.1\tau$, $t = 10\tau$ and $t = 1000\tau$) added for each transition.

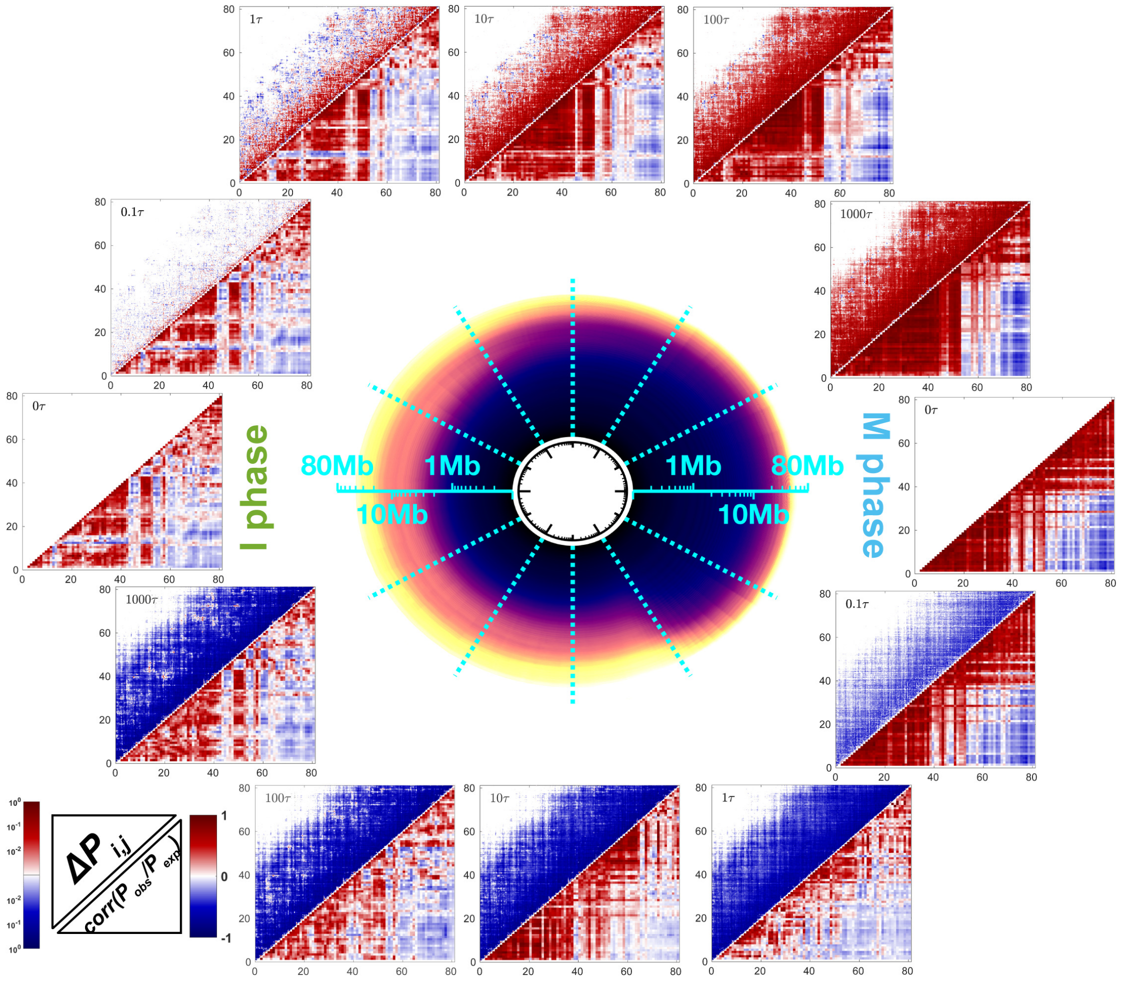


**FIG. S14:** Chromosome conformational transition during the cell cycle. The circle in the center is the same with that presented in Fig. 2 in the main text. At each matrix plot panel, the left-upper triangle is the difference of the contact probability $P_{i,j}(t)$ of the genomic locus pair $i$ and $j$ at time point $t$ to that at the initial ensembles with expression: $\Delta P_{i,j}(t) = P_{i,j}(t) - P_{i,j}(t=0)$; the right-lower triangle is the Pearson correlation matrix of the observed/expected contact probability map.

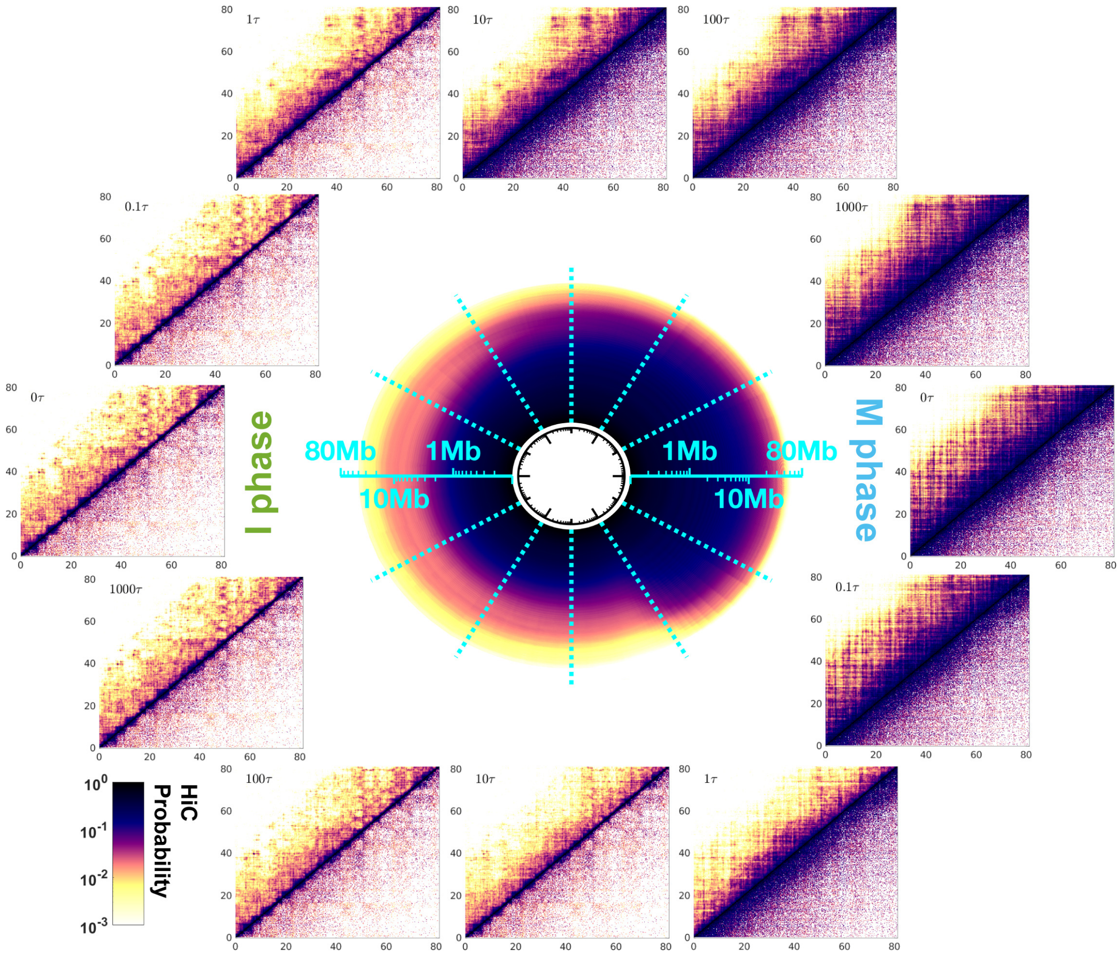


**FIG. S15:** Chromosome conformational transition during the cell cycle. The circle in the center is the same with that presented in Fig. 2 in the main text. At each matrix plot panel, the left-upper triangle is the contact probability $P_{i,j}(t)$ of the genomic locus pair $i$ and $j$ at time point $t$; the right-lower triangle is the Hi-C data. The I phase Hi-C data is used in the I→M transition, when $t = 0\tau$, $0.1\tau$ and $1\tau$, and in the M→I transition, when $t = 10\tau$, $100\tau$ and $1000\tau$. The M phase Hi-C data is used in the I→M transition, when $t = 10\tau$, $100\tau$ and $1000\tau$, and in the M→I transition, when $t = 0\tau$, $0.1\tau$ and $1\tau$.

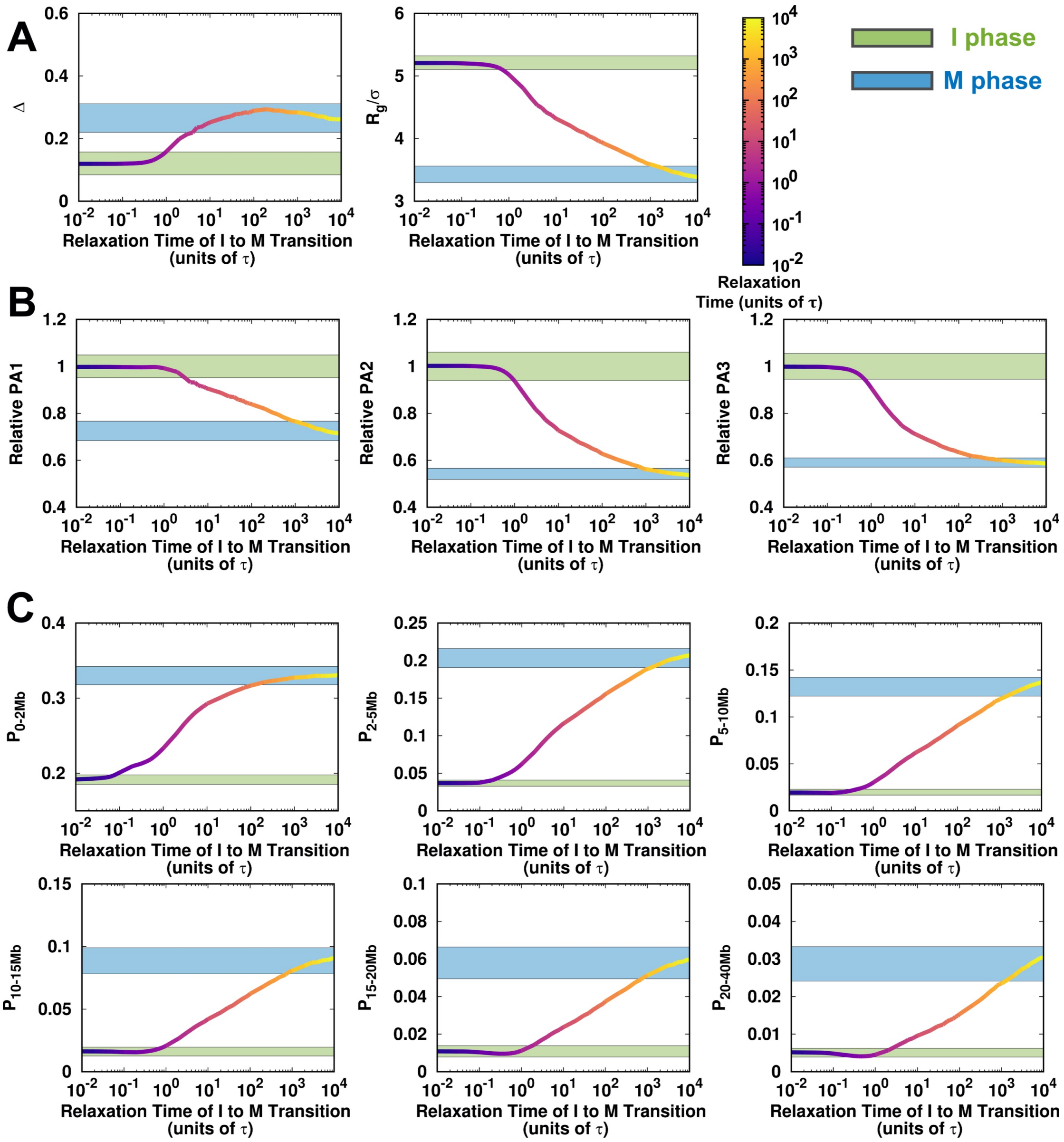


**FIG. S16:** The average pathways of the chromosome conformational changes during the I to M transition. The pathways are shown into three scenarios by projecting onto (A) shape quantities $\Delta$ and $R_g$, (B) extensions along PA1, PA2 and PA3, and (C) contact probability $P_l$ at various ranges $l$. The shadow ranges colored green and blue correspond to the values of the quantities obtained at the I and M phase. The widths of shadows represent the standard deviations of the corresponding quantities.

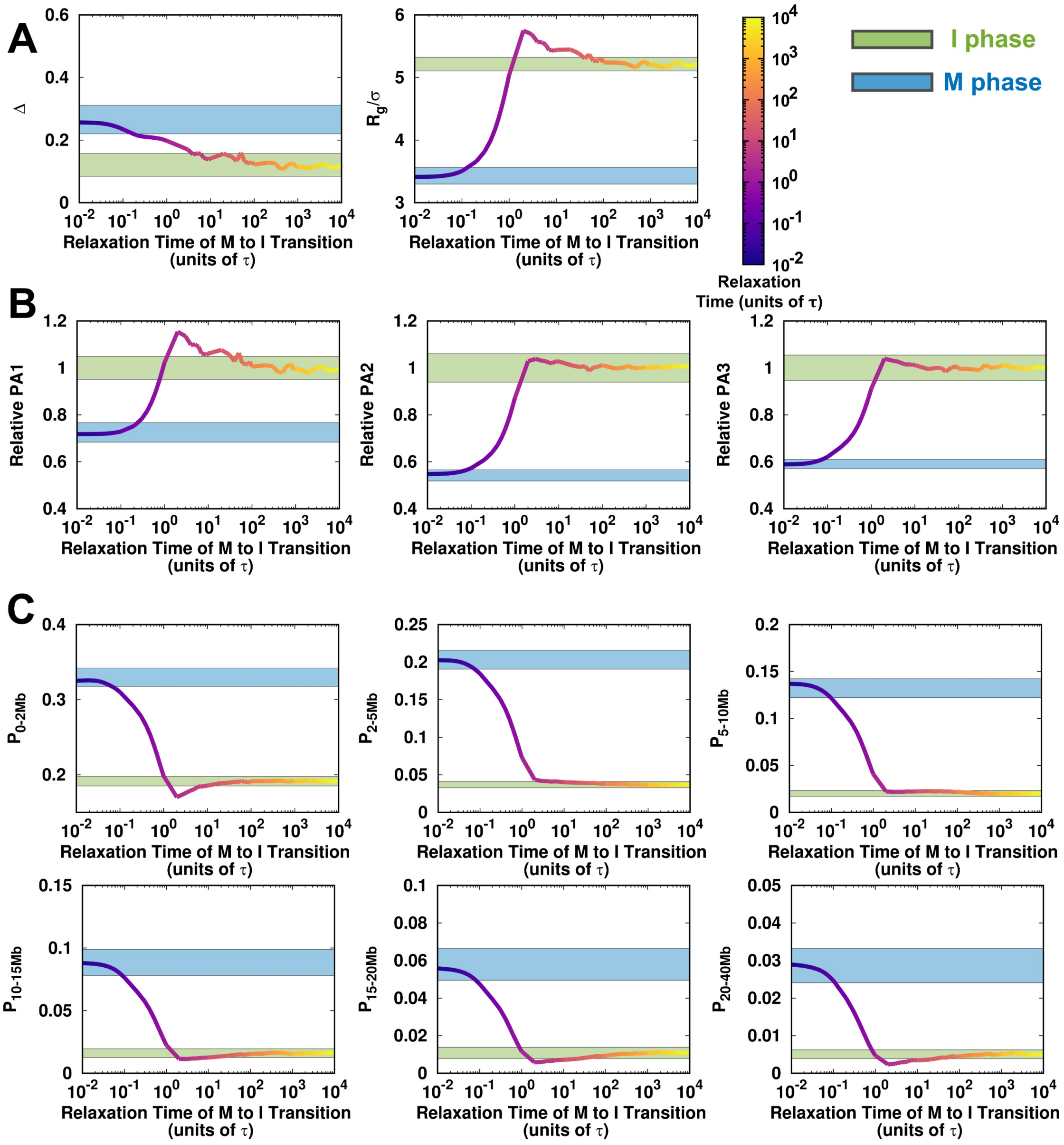


**FIG. S17:** The average pathways of chromosome structural changes during the M to I transition. Plots are the same with those in Fig. S16.

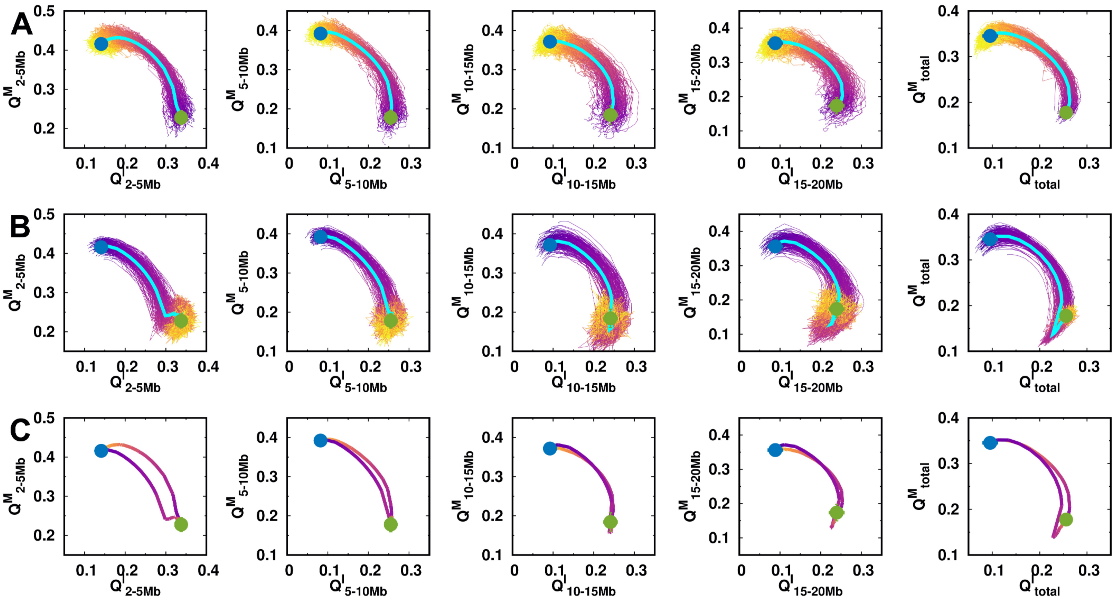


**FIG. S18:** The pathways projected onto the fraction of contact formation $Q$ during the cell cycle. (A) I→M transition. (B) M→I transition. (C) Average pathways for the two-directional transitions.

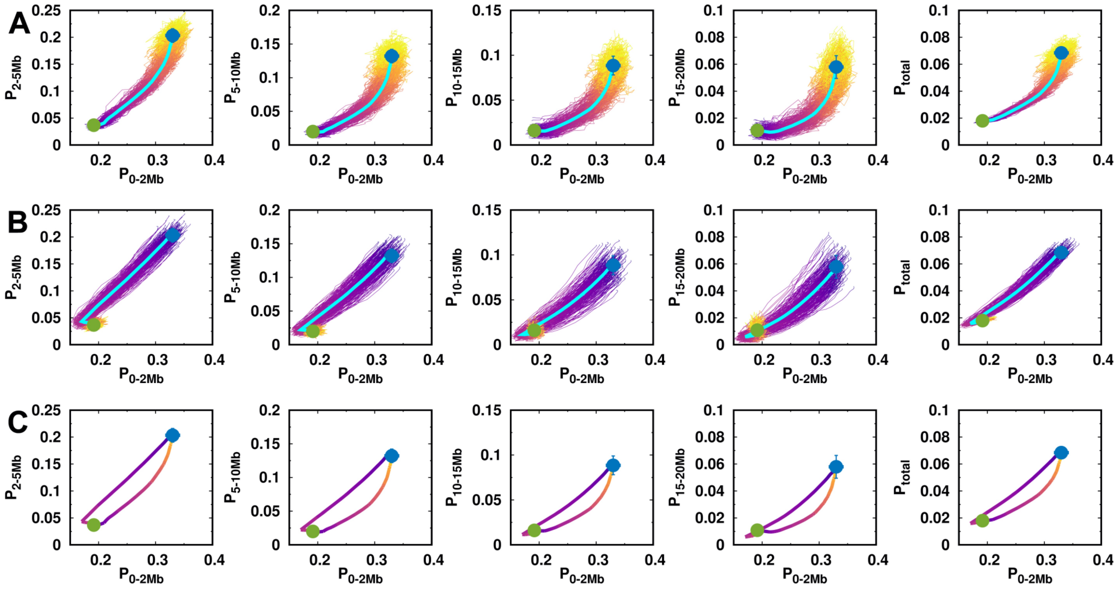


**FIG. S19:** The pathways projected onto the contact probability formation $P_l$ for various ranges during the cell cycle. (A) I→M transition. (B) M→I transition. (C) Average pathways for the two-directional transitions.

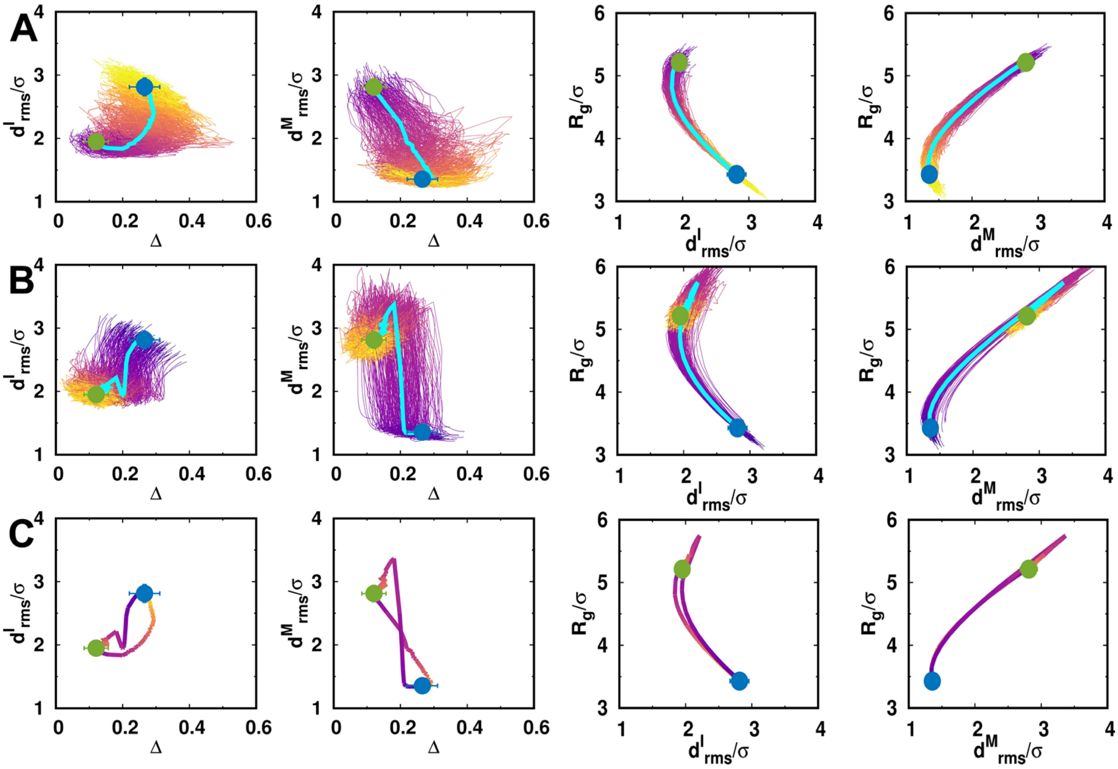


**FIG. S20:** The pathways projected onto $d_{rms}$, the aspheric shape parameter $\Delta$ and the radius of gyration $R_g$ during the cell cycle. (A) I→M transition. (B) M→I transition. (C) Average pathways for the two-directional transitions.

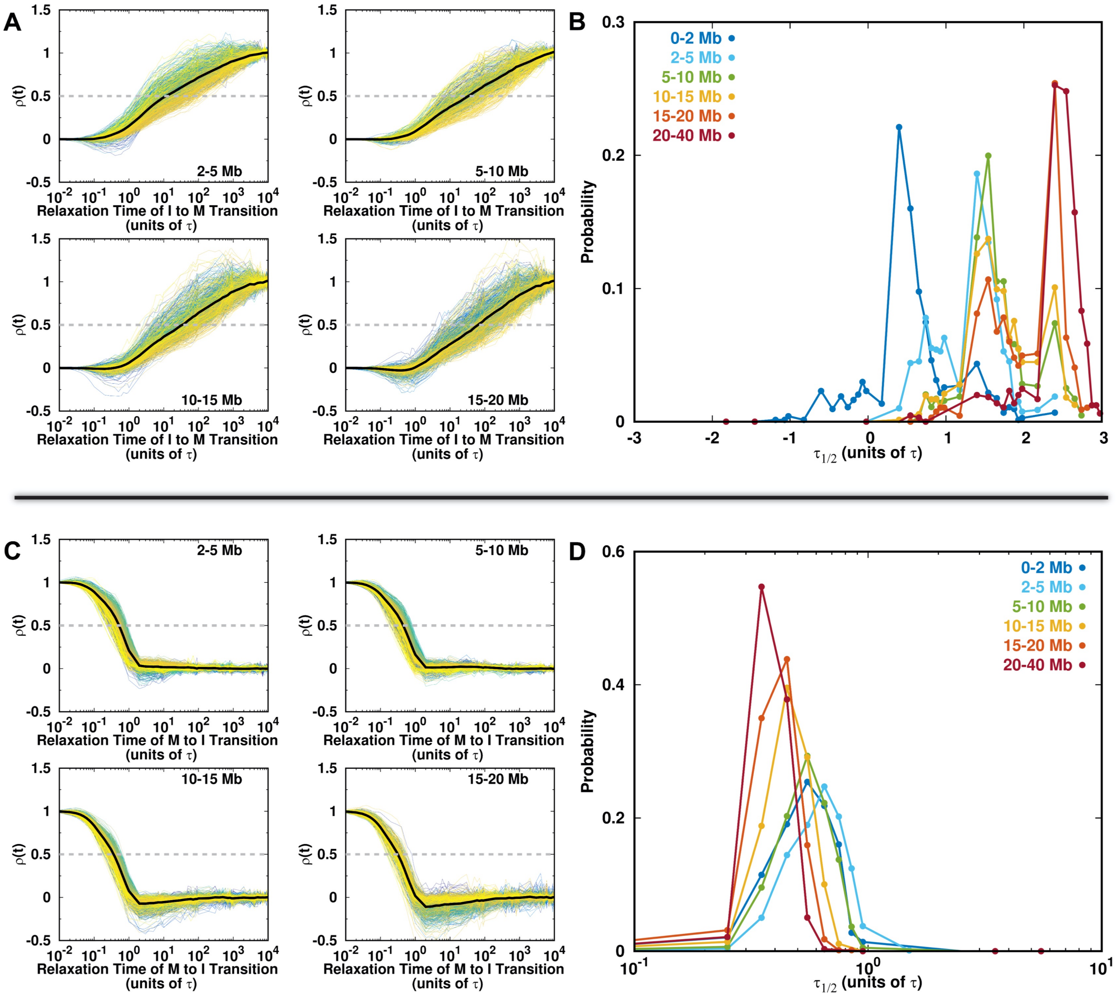


**FIG. S21:** Degree of the chromosome contact formation evolution during the cell cycle. (A) Different local and non-local contact formation evolutions for the transition from the I to M phase. (B) The half-life probability distributions at different genomic distances for the transition from the I to M phase. (B) and (D) the same with (A) and (C) but for the transition from the M to I phase.

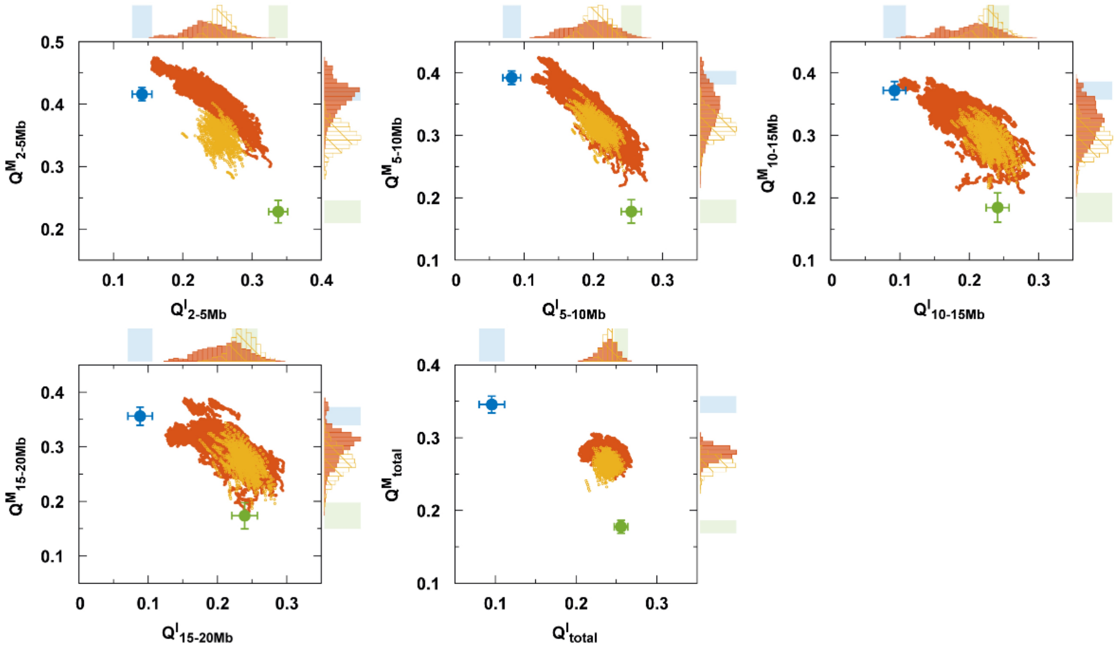


**FIG. S22:** Structural TS ensembles characterized by the fraction of contact $Q$ at different genomic distances.

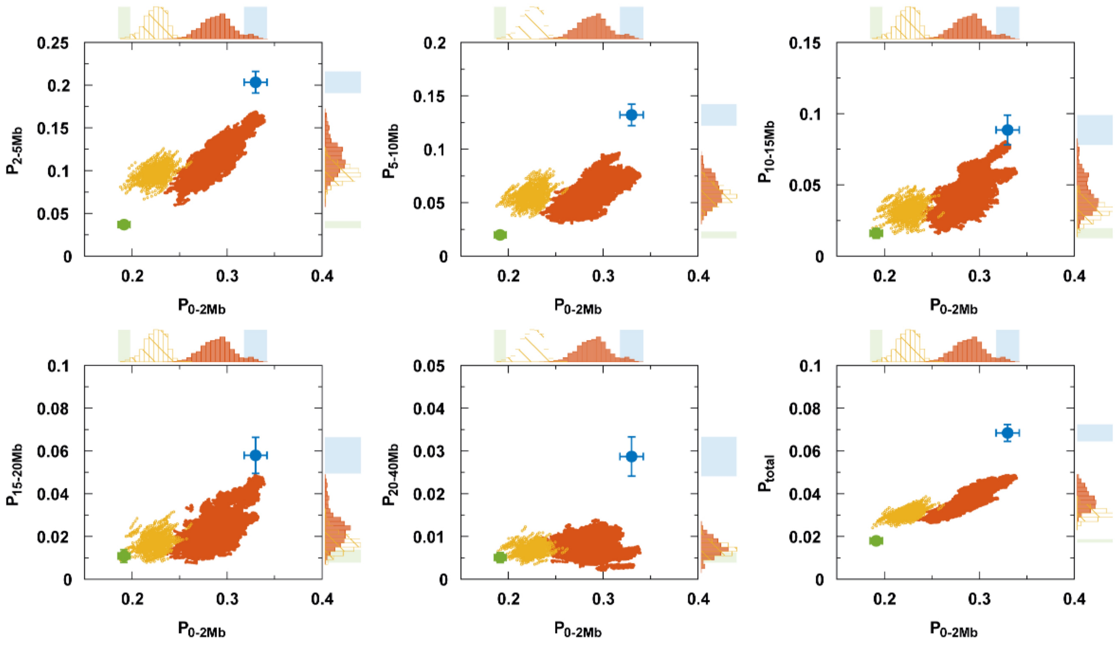


**FIG. S23:** Structural TS ensembles characterized by the contact probability $P$ at different genomic distances.
